# Characterising the effect of age and sex on post-transcriptional regulation in synovial joint tissues

**DOI:** 10.1101/2025.09.18.677035

**Authors:** Ufuk Ersoy, Sam Haldenby, Ioannis Kanakis, Julie Aspden, Mandy J Peffers, Simon R Tew

## Abstract

Age and sex are major risk factors for joint degeneration and disease susceptibility. While epigenetic and post-transcriptional mechanisms are known to be influenced by these factors, their effects on mRNA kinetics and configuration in musculoskeletal tissues remain poorly defined. To address this gap, we used an equine model to examine how age and sex impact mRNA stability in joint tissues. We measured global transcript half-life in primary chondrocytes from young and old female and young male horses using SLAM-seq. Polyadenylation patterns were additionally analysed in cartilage and synovium across age groups. Our findings demonstrated that age and sex exert distinct regulatory influences on post-transcriptional gene expression in joint tissues. Specifically, ageing alters polyadenylation site usage and transcript turnover in cartilage, even in absence of overt pathology, suggesting that molecular ageing may precede and predispose to joint degeneration. We also found that mean RNA half-life differed significantly between young females and males, with male chondrocytes showing greater transcript stability. This difference suggests that sex-specific regulatory mechanisms may influence RNA stability, which could contribute to differential susceptibility to joint degeneration. Together, these results point to age and sex as key drivers of post-transcriptional regulation with potential roles in shaping joint health trajectories.

**Graphical Abstract:** 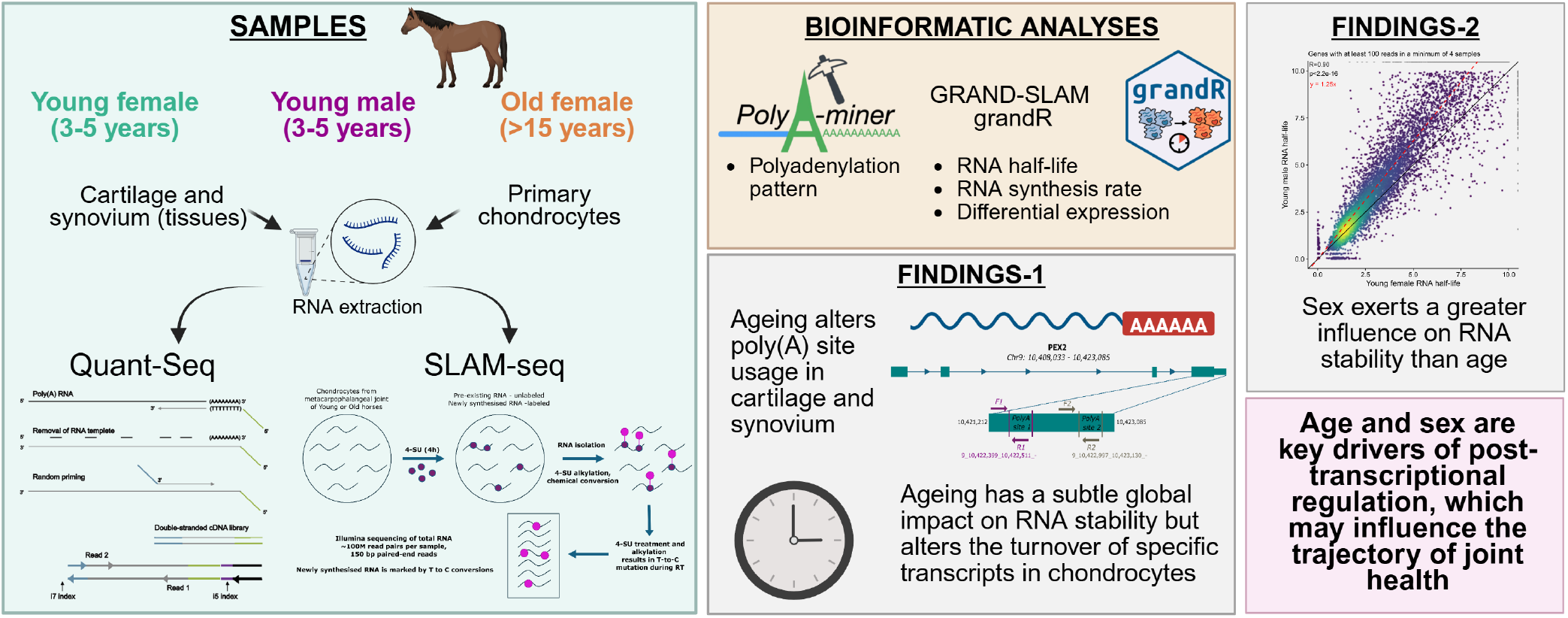

## Introduction

Ageing is a complex biological process, driven by a myriad of molecular and cellular processes which ultimately lead to a gradual decline of physiological function and an increased vulnerability to disease. It has become increasingly evident that the regulation of gene expression plays a pivotal role in orchestrating the effects of ageing, with changing patterns of transcript abundance identified in a range of ageing tissues (1–4). Furthermore, variations in epigenetic and post-transcriptional mechanistic processes have been associated with ageing (5–8). Key features of the latter include the kinetics of mRNA synthesis and turnover, as well as mRNA configuration, but these are poorly understood in synovial joint tissues in the ageing process.

Sex is a fundamental biological variable that influences physiology, disease susceptibility, and therapeutic responses (9,10). Accumulating evidence demonstrates widespread sex dimorphism in the prevalence and progression of age-related diseases. For example, osteoarthritis (OA) is more prevalent in females T after the age of 50, with women accounting for approximately 60% of global cases (11). Moreover, molecular mechanisms underlying ageing, including transcriptional regulation, are increasingly recognised to operate in a sex-dependent manner (12). However, the effects of sex on transcript kinetics remain largely unexplored, particularly in synovial joint tissues where sex-specific differences may contribute to age-related disease risk and responsiveness.

Transcript turnover rates are critical for modulating steady-state mRNA levels and shaping the responsiveness of genes to stimulation and repression (13–15). Widespread shifts in turnover rates of mRNAs are observed in cells exposed to stress or inflammatory stimuli and in pathological conditions, including cancer and neurodegeneration (16–22). In addition, polyadenylation plays a key role in mRNA stability, transport, and translation. Many transcripts undergo polyadenylation at multiple sites, and the selective inclusion or exclusion of RNA sequences through alternative polyadenylation (APA) serves as a key regulatory mechanism that influences transcript stability, kinetics, and localisation. This process of APA has been associated with a variety of diseases, including neurodegenerative disorders, metabolic syndrome, and genetic disorders (23,24). However, the effects of ageing and sex on mRNA turnover, transcript kinetics, and structure remain largely unexplored in synovial joint tissues, leaving significant gaps in our understanding of how transcript dynamics shape gene responsiveness across different physiological contexts.

Synovial joints are highly specialised biological environments bounded by either articular cartilage or the synovial lining/synovium (hereafter referred to as ‘synovium’) (25). The articular cartilage is an extracellular matrix-rich structural tissue containing a single cell type, the chondrocyte, which provides a low-friction, mechanically resilient surface to the bone within the joint (26). The synovium is a thin, membranous tissue covering the rest of the joint capsule and contains a heterogeneous population of fibroblasts, alongside immune cells such as macrophages, which are critical for the maintenance of inflammatory signalling within the joint (27). We have previously measured mRNA half-life in human articular chondrocytes and shown a propensity for shorter-lived transcripts in those isolated from osteoarthritic tissue (28). More recently, we have identified that OA can drive changes in polyadenylation site usage in a handful of chondrocyte gene transcripts, but that the effect at a transcriptomic scale was modest (29).

A challenge of characterising cellular mechanisms in human OA is that, while diseased tissue can be obtained following joint arthroplasty, the supply of healthy tissue is limited and often comes from individuals with different characteristics, particularly age and sex. This limitation is especially critical for kinetic analyses, which require fresh material. In our previous half-life study, control tissue could only be sourced from younger individuals, which constrained our ability to assess the influence of these variables (28). To address this, we aimed to understand the extent to which age and sex influence post-transcriptional regulatory mechanisms in cartilage and synovium. We therefore employed an equine model system, benefiting from animals with well-documented records that allow for age- and sex-matched sample selection. The horse is widely recognised as a suitable model for studying human joint biology and osteoarthritis due to the comparable thickness, structure, and mechanical properties of equine articular cartilage to those of humans (30,31). The equine model also offers several advantages for ageing research, including the ability to control for environmental and lifestyle factors such as diet and smoking, which can confound human studies through epigenetic effects. We assessed global transcript half-life in primary chondrocytes isolated from young female and male, as well as old female animals, using metabolic RNA labelling to evaluate the effects of both age and sex. Complementary analyses of polyadenylation patterns were conducted in cartilage and synovium, but only across age groups. We found that age and sex exert distinct effects on post-transcriptional gene regulation in joint tissues. Ageing alters both polyadenylation site usage and transcript turnover in cartilage, while sex has a pronounced influence on mRNA stability in chondrocytes, potentially contributing to differences in susceptibility to joint degeneration. These results highlight age and sex as key regulators of RNA dynamics with important implications for joint health.

## Materials and Method

### Collection of tissue

Equine lower limbs were obtained at the time of slaughter of young (3-5 years) or old animals (*>*15 years) from the F Drury and Sons abattoir, Swindon, UK. As the samples were collected as a byproduct of the agricultural industry, ethical approval was not required for the study. The joints were dissected, photographed and scored using the Osteoarthritis Research Society International equine macroscopic grading approach (32), with any showing evidence of pathology excluded from the study (Supplementary Table 1). For the analysis of APA, articular cartilage, harvested from the metacarpus, proximal phalanx, and sesamoids, or synovium was dissected from the metacarpophalangeal joint, snap frozen in liquid nitrogen, and stored at ™80°C(33). For transcript kinetic analysis, multiple pieces of articular cartilage from the same anatomical regions were dissected and processed for primary cell culture, as previously described (34).

### QuantSeq: sample preparation and APA analysis

Frozen tissue samples from young and old horses were homogenised using a Braun Mikrodismembrator and extracted in TRIzol (Thermo Fisher Scientific, Waltham, MA, USA, #15596018). Following the addition of chloroform and phase separation, the aqueous layer was removed and processed through RNeasy mini kits (Qiagen, Manchester, UK, #74104), including on-column DNase digestion. QuantSeq-Reverse library preparation and NextSeq 75 cycle High Output sequencing with V2chemistry was performed by Lexogen (Vienna, Austria).

### Identification of polyadenylation site and age-associated APA events

Polyadenylation sites were identified and their usage compared as a result of age or tissue type. QuantSeq reverse reads were analysed using PolyAMiner (35). An initial run was conducted with ‘-expNovel 1’ to explore novel APA sites. De novo APA sites were extracted and peaks with a maximum size of 1 kb were assigned to genes within 5 kb, respecting strand orientation, using BEDTools ‘closest’ (36). Filtered peaks associated with genes were then used as input as polyA annotations (option ‘-pa’) to PolyAMiner.

### Quantification of APA site usage with qPCR

Snap-frozen cartilage or synovium samples (n = 5-6) were ground in liquid nitrogen using a mortar and pestle. Total RNA was extracted using TRIzol and purified with RNeasy Mini Kits as previously described (37,38). cDNA synthesis was performed using the High-Capacity cDNA Reverse Transcription Kit (Applied Biosystems, Warrington, UK, #4368814), primed with 2 *µ*M oligo(dT) to selectively target polyadenylated transcripts.

Quantitative PCR was conducted on a Roche LightCycler 96 using Takyon ROX SYBR MasterMix (Eurogentec, Seraing, Belgium, #UF-FSMT-B0701). Isoform-specific primers were designed to distinguish transcripts based on polyA site usage. Primer efficiency and validation were carried out as previously described using genomic DNA (29). Where there are two significantly differently used APA sites located in the last exon, the expression levels were first normalised to GAPDH. The normalised Ct values for each isoform were then used to calculate Δ*C*_t_ (Ctdistal - Ctproximal). ΔΔ*C*_t_ values were derived by subtracting the average Δ*C*_t_ of the control group from each sample’s Δ*C*_t_. The relative polyA site usage was expressed as fold change using 2^*−*ΔΔ*C*t^ method. For significantly differentially used intronic APA sites, site-specific transcript abundance was normalised to GAPDH expression, and fold change was calculated using the standard 2^*−*ΔΔ*C*t^ method (37). Primer sequences are listed in Supplementary Table 2.

### Western blotting

Total protein was extracted from a portion of the ground synovium using RIPA buffer (Sigma, Poole, UK, # R0278) supplemented with Pierce Protease and Phosphatase Inhibitor Mini Tablets, EDTA-free (Thermo Fisher Scientific, Waltham, MA, USA, #A32961). The samples were homogenised and sonicated twice for 30 seconds, then centrifuged at 14,000Xg for 10 minutes at 4°C. The supernatant was collected, and total protein concentration was determined using the BCA Protein Assay (Merck, Millipore, Burlington, Massachusetts, US, #71285-M).

30 *µ*g of protein were separated on a NuPAG− 4–12% Bis-Tris Mini Protein Gel (Invitrogen, Renfrewshire, UK, #NP0321BOX) and transferred to a polyvinylidene fluoride (PVDF) membrane (Roche, Basel, Switzerland, #3010040001). Ponceau S staining (Sigma, Poole, UK, #P7170) was used to monitor consistent loading. The membrane was blocked with 5% bovine serum albumin (BSA) in Tris-buffered saline (TBS) for 1 hour at room temperature (RT). The membrane was incubated overnight at 4°C with anti-NRG1 antibody (1:600, Proteintech, Rosemont, IL, USA, #66492-1), followed by incubation with a secondary anti-mouse LI-COR antibody at RT. The bands were visualised using the LI-COR Odyssey CLx imaging system (LI-COR, Bad Homburg, Germany). Band densities were analysed with ImageJ and normalised to Ponceau staining (39).

### Cell culture and metabolic RNA labelling with 4-thiouridine (4sU)

Dissected articular cartilage was finely cut into 1-3 mm pieces with a scalpel, treated with 1x trypsin in serum free cell medium containing 100 units/ml penicillin-streptomycin (Pen/Strep; Gibco, #15140122), and then digested in a 0.08% collagenase type II solution (Worthington Biochemicals, Lakewood, NJ, USA, #LS004176) at 37°C overnight. Once strained and washed, cells were seeded at 100,000 cells*/*cm^2^ in tissue culture plastic and cultured in complete medium: Dulbecco’s Modified Eagle Medium (DMEM; Thermo Fisher Scientific, #31885023) with 10% foetal bovine serum (FBS; Sigma, #F7524) and 100 units/ml Pen/Strep.

Following chondrocyte isolation and culture, cells were incubated for 2 days. The medium was changed, and 4-thiouridine (4sU; Sigma-Aldrich, Gillingham, Dorset, UK, #T4509) was added to a final concentration of 500 *µ*M for 4 hours at 37°C. After 4sU treatment, cells were washed twice with prewarmed phosphate-buffered saline (PBS). The total RNA was then harvested using TRIzol reagent, collected in 1.5 ml Eppendorf tubes (Eppendorf, Hamburg, Germany) and stored at *−*80°C.

### SLAM-Seq: chemical conversion and library preparation

Chemical conversion of the labelled RNA was performed following the protocol by Herzog et al. (40). Briefly, total RNA was isolated from TRIzol extracts using isopropanol precipitation with the addition of 0.1 mM dithiothreitol (DTT; Thermo Fisher Scientific, Waltham, MA, USA, #A39255). After precipitation, RNA was washed in 1 mM DTT in 75% ethanol and resuspended in RNase-free water. A total of 5 *µ*g RNA was treated with iodoacetamide (IAA; Thermo Fisher Scientific, Waltham, MA, USA, #122271000) at a final concentration of 10 mM for 15 minutes at 50°C. Alkylation was stopped by quenching the reaction with 1 mM DTT (final concentration), and then the total RNA was ethanol precipitated and subjected to library preparation as below.

The preparation of Illumina RNA libraries from total equine RNA samples was performed using the NEBNext polyA selection and Ultra II Directional RNA library prep kits (New England Biolabs, UK, #E7760S), as previously described (3). Libraries were multiplexed and sequenced on half a lane of a NovaSeq X Plus 25B flow cell. A minimum of 78.1 million paired-end reads (2 × 150 bp) were sequenced per replicate (Figure 4a). Sequencing was performed at the Centre for Genomic Research, University of Liverpool (https://www.liverpool.ac.uk/genomic-research/).

### SLAM-Seq data processing and bioinformatic analyses

GRAND-SLAM (41) and grandR (42) were used to estimate the half-life of RNA (hours), the synthesis rate (TPM/hour), and differential expression. The raw FASTQ files were trimmed to remove Illumina adapter sequences with Cutadapt version 4.5. Reads shorter than 15 bp were discarded after trimming. Only read pairs were retained; if one read of a pair was removed, its mate was also discarded. The reads were aligned to the Equus caballus EquCab3.0 primary assembly (Ensembl release 114) using STAR (version 2.5.3) with the following parameters: –outFilterMismatchNmax 40, –outFilterScoreMinOverLread 0.2, –outFilterMatchNminOverLread 0.2, –alignEndsType Extend5pOfReads12, and –outSAMattributes nM MD NH. The GEDI toolkit was utilised to merge and convert BAM files into a CIT file (43) and then processed using GRAND-SLAM (version 2.0.7) to generate new-to-total RNA values (NTR) and read counts. RNA synthesis rates and half-lives were estimated with the grandR using binomial distributions for reads from labelled and unlabelled RNAs.

grandR (version 0.2.6) was used to generate the Principal Component Analysis (PCA) plot, scatter plots, progressive time course plots, volcano plot, and MA plot in R (version 4.4.1). RNA half-life distributions were visualised using ggplot2 (version 3.5.2). Empirical cumulative distribution functions (ECDFs) were also generated with ggplot2 to compare half-life distributions between conditions on a log-transformed scale. Additionally, scatter plots of RNA half-life versus transcript abundance (TPM) were created using ggplot2 with genes having |*logfoldchange*(*LFC*)| *>* 0.5 and a Q value *<* 0.05. A list of genes filtered by RNA half-life changes was input into ShinyGO (version 0.8) to perform Gene Ontology (GO) analysis (44).

Differential expression analysis was performed using the PairwiseDESeq2 function from the grandR. RNA half-life categories were defined based on mean half-life values per gene across all samples, grouped into five intervals: 0–2h, 2–4h, 4–8h, 8–10h, and *>*10h. LFCs were then compared across these categories using boxplots generated with ggplot2. A heatmap of differentially expressed genes was generated using the ComplexHeatmap package (v2.22.0) in R. Gene set enrichment analysis (GSEA) was performed on differentially expressed genes using the AnalyzeGeneSets function in grandR, focusing on GO and Kyoto Encyclopaedia of Genes and Genomes (KEGG) terms for Equus caballus. The top enriched pathways were visualised with a ridge plot generated with clusterProfiler (version 4.14.6).

### Statistical analyses

Genes significant for differential APA were identified with PolyA-miner by quantifying changes using a likelihood ratio test (LRT) statistic, derived from beta-distributed co-clustering frequencies, which is approximated as a chi-squared distribution for statistical significance (35). In the SLAM-seq analysis, GRAND-SLAM (41) and grandR (42) were used to estimate RNA half-life, synthesis rate, and differential expression. The t-test for qRT-PCR analysis and western blot was computed using the rstatix package (version 0.7.2). Differential expression and PCA were computed as described above. All Pearson correlation coefficients were calculated using the cor function of R (version 4.4.1).

## Results

### Altered polyadenylation patterns are associated with age in joint tissues

The horse metacarpophalangeal joint is subjected to substantial load throughout an animal’s life and is susceptible to age-associated degenerative diseases such as OA (45). To investigate how age and sex influence mRNA stability in joint tissues, we examined samples from male and female horses aged 3–5 years (representing young adults) and from females over 15 years of age (representing old age in humans) (Figure 1a).

**Fig. 1.**
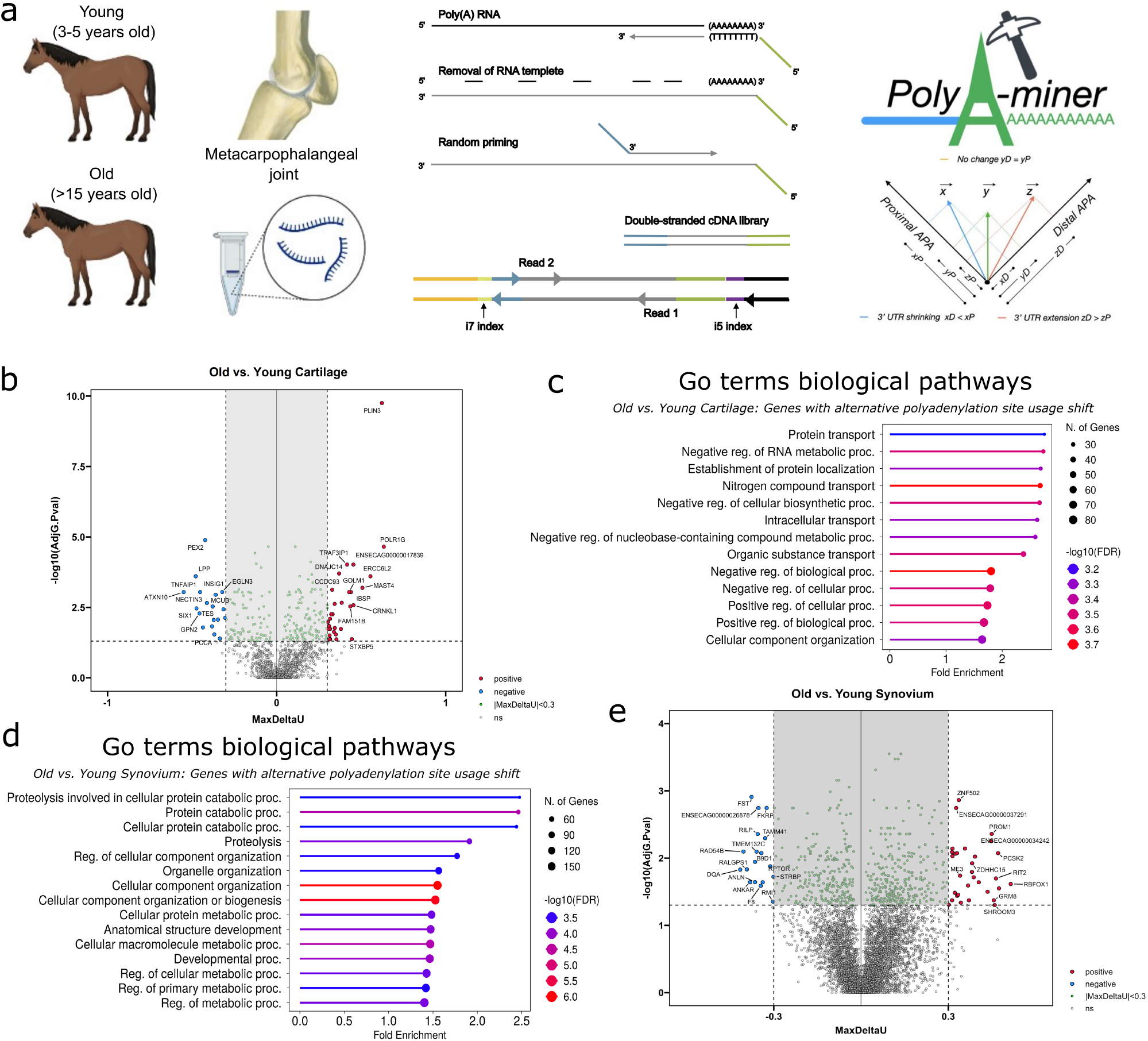
Altered polyadenylation patterns are associated with age in joint tissues (a) Schematic of the experimental workflow for detecting differential alternative polyadenylation (APA) site usage. Total RNA was isolated from cartilage and synovium of the metacarpophalangeal joints of young (3–5 years old) and old (*>*15 years old) horses. Libraries were prepared using QuantSeq-Reverse and sequenced on an Illumina NextSeq platform. APA site switching events were identified using the PolyA-miner software (35). Volcano plot indicating the maximum change in polyadenylation site usage (MaxDeltaU) for genes in (b) cartilages and (e) synovium, comparing samples from aged and young female horses. Genes with significantly increased site usage in old tissue are highlighted in red; those with significantly decreased usage are highlighted in blue. Gene Ontology (GO) biological process enrichment analysis of genes showing significant shifts in APA site usage in (c) cartilage and (d) synovium. Circle colour represents the –log_10_(FDR) associated with pathway enrichment, and circle size reflects the number of genes associated with each GO term.

We sought to determine the effect of age on polyadenylation site usage in joint tissues. Using total RNA purified from articular cartilage or synovium, we performed QuantSeq-Rev 3′ end RNA sequencing to precisely map polyadenylation sites. This method sequences from the 3′ end of transcripts, enabling accurate identification of polyadenylation events. We then utilised PolyAminer software and found evidence that age affected polyadenylation site usage in both cartilage and synovium (Supplementary Tables 3-6). In the articular cartilage, 243 genes exhibited evidence of APA between at least two sites (Figure 1b and Supplementary Figure 1a). Using gene enrichment analysis, the functionality of these genes was shown to be linked to a range of cellular transport processes, including protein transport, negative regulation of RNA metabolic processes, and establishment of protein localisation (Figure 1c). In the synovium, 590 genes exhibited differential usage of APA sites between young and old individuals. Despite this, the impact of ageing on APA site selection in this tissue was relatively modest (Figure 1e and Supplementary Figure 1b). Genes exhibiting differential APA usage in the synovium were associated with a range of catabolic, metabolic, and cellular organisation processes (Figure 1d).

To validate the polyadenylation data generated by the QuantSeq Reverse analysis, we designed qRT-PCR assays to assess the ratios of polyadenylated isoforms for a selection of genes in an independent cohort of young and old equine samples, comprising the same animals analysed by SLAM-seq in this study (Supplementary Table 1). For articular cartilage, we examined the Perilipin-3 (PLIN3) and Peroxisomal biogenesis factor 2 (PEX2) genes, which each demonstrate altered 3′UTR length as a result of the APA events. Transcriptomic analysis identified PLIN3 transcripts with an extension of 117 nt to the 3′UTR in young samples, whilst use of this polyadenylation site was barely detected in old samples (Figure 2a). qPCR supported the differential use of this site (Figure 2a). In contrast, PEX2 transcripts in older samples predominantly utilised a polyadenylation site that resulted in a 486 nt extension of the 3′UTR. PCR analysis in an independent cohort further confirmed the age-associated differential usage of this site (Figure 2b). In the synovium, we examined RNA-binding protein RBFOX1 and Prominin 1 (PROM1), as they exhibited the most significant differential APA usage, within the 3′UTR and intronic regions, in old samples compared to young. qRT-PCR primers showed that the relative expression patterns were consistent with those observed in the QuantSeq Reverse analysis; however, no statistically significant changes were detected (Figure 2c and 2d). This may be attributed to the cellular heterogeneity of the synovium. These findings reinforce that APA contributes to transcriptomic remodelling of joint tissues with age.

**Fig. 2.**
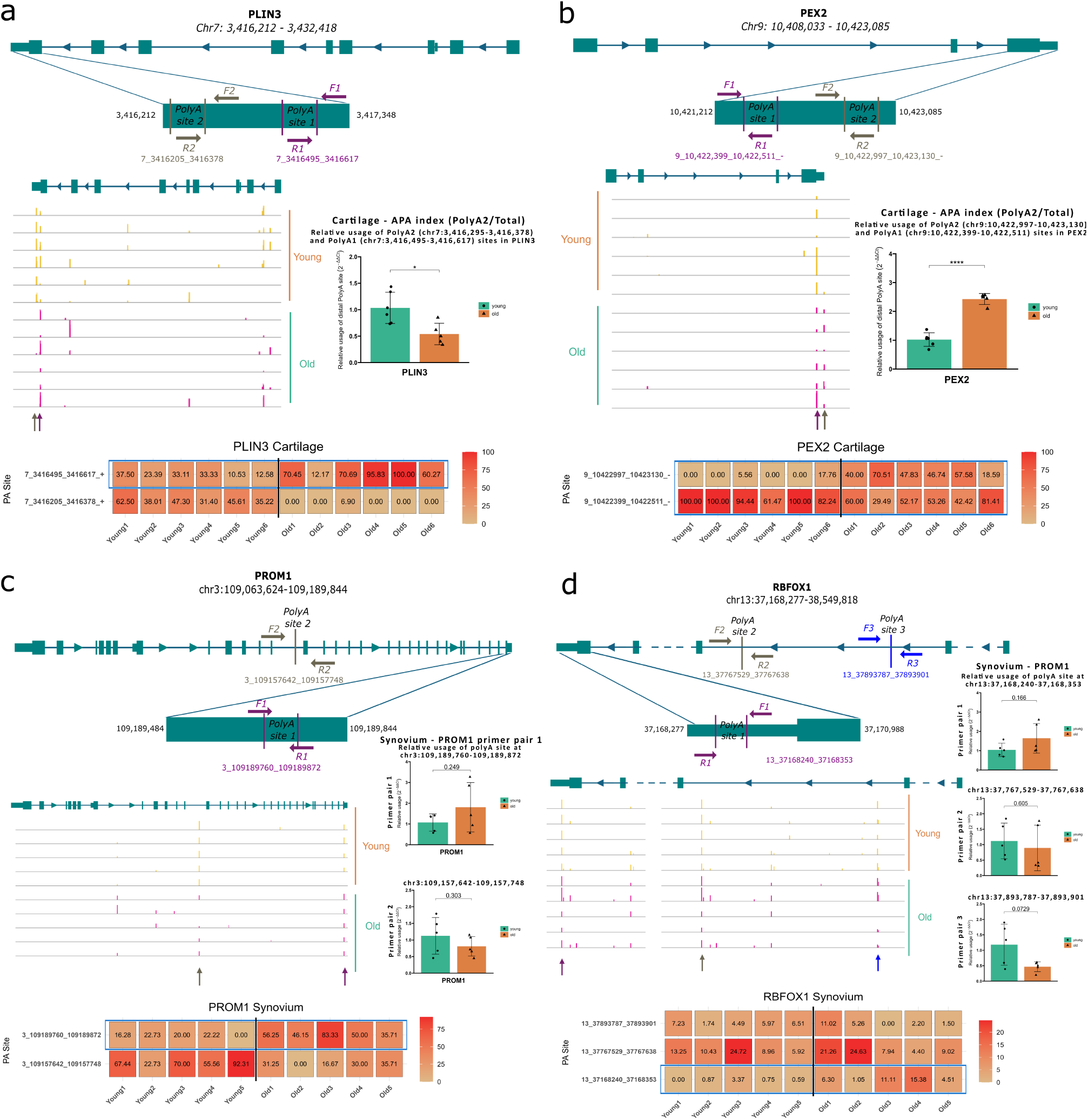
qPCR validation of QuantSeq-identified alternative polyadenylation (APA) changes in PLIN3 and PEX2 in cartilage, and PROM1 and RBFOX1 in synovium. (a) APA analysis of the PLIN3 gene in cartilage. The top panel shows the genomic organisation of PLIN3 on chromosome 7 (chr7:3,416,212–3,432,418) with two annotated polyadenylation sites: proximal (PolyA1, chr7:3,416,495–3,416,617) and distal (PolyA2, chr7:3,416,205–3,416,378). Primer pairs F1/R1 and F2/R2 flank PolyA1 and PolyA2, respectively. The middle panel shows QuantSeq read density tracks in cartilage from young (orange) and old (teal) samples. The right panel shows qPCR quantification of PolyA site usage, with PolyA1 primers amplifying two isoforms and PolyA2 primers specific to distal site transcripts. Ct values were normalised to GAPDH and analysed using the 2^*−*ΔΔ*C*_t_^ method (mean *±* SEM, n=5–6), revealing a significant age-related reduction in distal site usage. (b) APA analysis of the PEX2 gene in cartilage. The top panel shows the genomic organisation of PEX2 on chromosome 9 (chr9:10,408,033–10,423,085) with two annotated polyadenylation sites: proximal (PolyA1, chr9:10,422,997–10,423,130) and distal (PolyA2, chr9:10,422,399–10,422,511). Primer pairs F1/R1 and F2/R2 flank PolyA1 and PolyA2, respectively. The middle panel shows QuantSeq read density tracks in cartilage from young (orange) and old (teal) donors. The right panel shows qPCR quantification of PolyA site usage, with PolyA1 primers amplifying two isoforms and PolyA2 primers specific to distal site transcripts. Ct values were normalised to GAPDH and analysed using the 2^*−*ΔΔ*C*^t method (mean *±* SEM, n=5–6), revealing a significant age-related increase in distal site usage. (c) APA analysis of the PROM1 gene in synovium. The top panel shows the genomic organisation of PROM1 on chromosome 3 (chr3:109,063,624–109,189,844) with two annotated polyadenylation sites: a distal exonic site (PolyA1, chr3:109,189,760–109,189,872) and a proximal intronic site (PolyA2, chr3:109,157,642–109,157,748). Primer pairs F1/R1 and F2/R2 flank PolyA1 and PolyA2, respectively. The middle panel shows QuantSeq read density tracks in synovium from young (orange) and old (teal) donors. The right panels show qPCR measurements of site-specific transcript abundance, normalised to GAPDH expression. Data are presented as mean *±* SEM (n=5), with no statistically significant age-related differences detected at either site. (d) APA analysis of the RBFOX1 gene in synovium. The top panel shows the genomic organisation of RBFOX1 on chromosome 13 (chr13:37,168,277–38,549,818) with three annotated polyadenylation sites: PolyA1 (chr13:37,168,240–37,168,353), PolyA2 (chr13:37,767,529–37,767,638), and PolyA3 (chr13:37,893,787–37,893,901). Primer pairs F1/R1, F2/R2, and F3/R3 flank PolyA1, PolyA2, and PolyA3, respectively. The middle panel shows QuantSeq read density tracks in synovium from young (orange) and old (teal) donors. The right panels show qPCR measurements of site-specific transcript abundance, normalised to GAPDH expression. Data are presented as mean *±* SEM (n=5), with no statistically significant age-related differences detected at any site.

### Differences in cell type have a larger contribution to differing APA usage than age

We analysed APA data from synovium and cartilage to identify common age-associated changes. From over 800 genes that exhibited APA, only 17 were shared between the two tissue types (Figure 3a), and among these, neuregulin 1 (NRG1) exhibited APA switching change between young and old (Figure 3b). The NRG1 polyadenylation site resides in intron 1, and examination of this feature in the genome browser revealed two corresponding expressed sequence tags (ESTs), CD472152 and CD535840. BLAST analysis of these ESTs revealed over 92% similarity to eukaryotic translation elongation factor 1 alpha 1 (EEF1A1) in human and mouse. Given the extensive distance from the 3’ end of the exon 1 to this regulated polyadenylation site (*>*100 kb), it is more likely to represent a separate intronic transcript, rather than a transcriptional run through from the first exon. Consistent with this, analysis of protein extracts from synovium did not demonstrate altered levels of NRG1 protein abundance between young and old samples, and no evidence of any alternatively sized protein isoforms (Supplemental Figure 1c). To investigate further, we used PCR to examine transcripts derived from NRG1 intron 1 in an independent set of equine samples. PCR designed to detect total transcript levels (Figure 3e), we were able to detect transcript expression and observed that expression tended to increase in older samples, particularly in cartilage, aligning with the original QuantSeq Reverse data (Figure 3f). Use of a primer pair closer to exon 1 did not identify transcripts within these samples, indicating that the polyadenylation site is associated with an independently transcribed intronic RNA, rather than running on from NRG1 exon 1 (Figure 3e). Together, these findings suggest that this region may contain a retrotransposed EEF1A1-like pseudogene fragment that may be elevated in aged tissue.

**Fig. 3.**
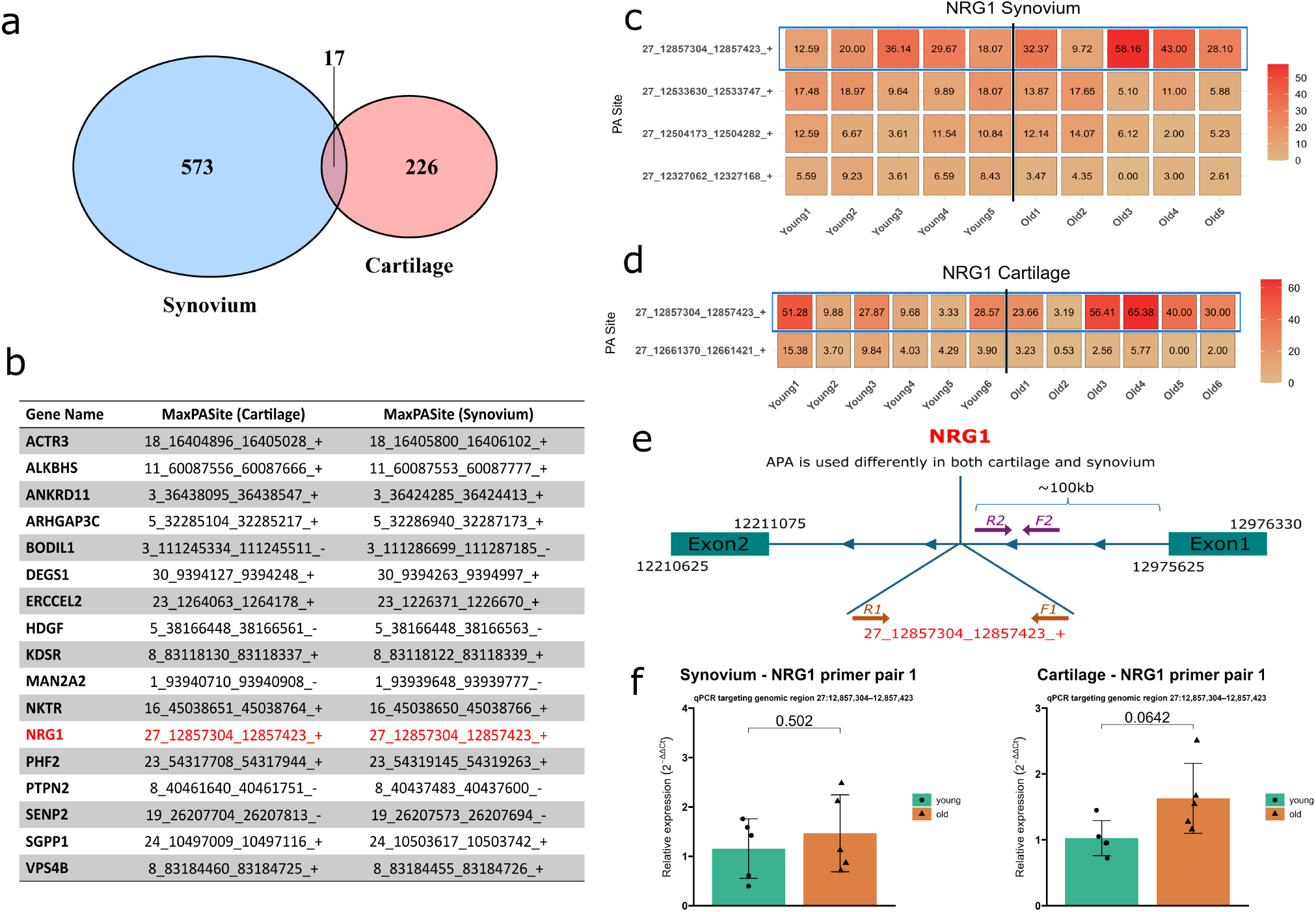
Differences in cell type have a larger contribution to differing APA usage than age (a) Venn diagram showing the number of genes with differentially used APA sites in synovium and cartilage. Synovium contains more genes with APA changes, with 17 genes shared between both tissues. (b) Table listing these 17 genes with differential APA site usage detected in both synovium and cartilage. Only NRG1 shows consistent polyadenylation site usage across both tissues. (c, d) Normalised APA proportion matrices of the NRG1 in (c) synovium and (d) cartilage. Usage of the intronic APA site at 27 12857304 12857423 + is annotated in blue. n = 5-6. (e) qPCR assay strategy: Primer pair 1 was designed to determine whether the intronic APA site at 27 12857304 12857423 + is used to transcribe a small non-coding RNA. Primer pair 2 was designed to assess whether the usage of the same APA site results in a truncated mRNA. (f) qPCR analysis of NRG1 using primer pair 1 was performed on synovium and cartilage samples from young and old horses. n = 5. Statistical significance was determined using an unpaired t-test, with p *<* 0.05 considered significant.

**Fig. 4.**
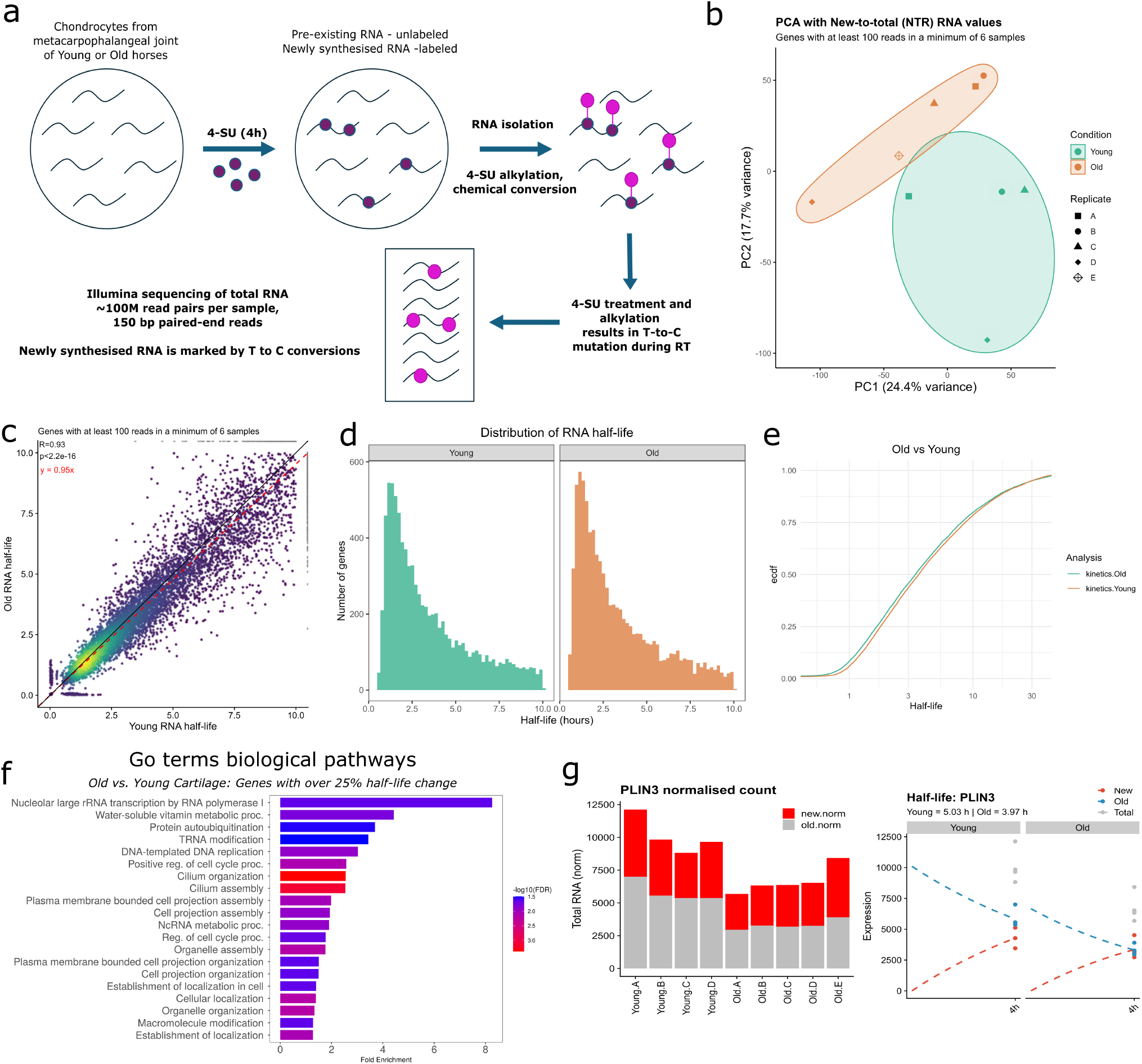
SLAM-Seq analysis of RNA half-lives. (a) Schematic illustration of the thiol (SH)-linked alkylation for the metabolic sequencing of RNA (SLAM-seq) approach (40). SLAM-seq was performed in chondrocytes isolated from the metacarpophalangeal joint of young and old horses. Newly synthesised RNA was labelled with 4-thiouridine (4sU) for 4 hours. RNA was isolated and treated with Iodoacetamide (IAA). n = 5. Libraries were prepared and sequenced on an Illumina NextSeq platform. (b) PCA analysis with new-to-total (NTR) RNA values. Genes included had 100 reads in at least 6 samples. (c) Scatter plot showing the relationship between Old Half-life and Young Half-life for genes with at least 100 reads in a minimum of 6 samples. A linear regression line (black) is fitted to the data, with the equation y=0.95x (red), indicating minimal changes in half-lives. Point density is represented by a colour gradient from blue (low density) to yellow (high density). (d) Distribution of RNA half-lives measured in chondrocytes from young and old horses by SLAM-seq. (e) Empirical cumulative distribution function (ECDF) of RNA half-life for chondrocytes from young and old horses. The ECDF illustrates the proportion of genes with half-lives below a given threshold across both age groups. (f) Gene Ontology (GO) biological process enrichment analysis of genes exhibiting *>*25% change in RNA half-life in old cartilage samples compared to young. Bar length indicates fold enrichment, and bar colour represents –log_10_(FDR) values associated with pathway enrichment, ranging from blue (lower significance) to red (higher significance). (g) Normalised total RNA counts for PLIN3 in young and old samples. Bars represent newly synthesised (red) and pre-existing (grey) RNA components. Progressive labelling plots demonstrating expression dynamics of PLIN3 in young and old samples. Plots show newly synthesised (red dashed) and pre-existing (blue dashed), and total (grey dots) RNA, with estimated half-lives of 5.03 hours (young) and 3.97 hours (old).

### SLAM-seq analysis of RNA half-lives in chondrocytes

We next investigated how age and sex affected mRNA kinetics in chondrocytes isolated from young female and male horses, as well as from older females. We focused on this cell type due to its critical role in articular cartilage maintenance and the relative ease of isolating appropriate cell numbers from the equine metacarpophalangeal joint to permit prompt analysis and avoid cell dedifferentiation. SLAM-seq was used to measure the levels of detection of newly synthesised and pre-existing RNA within one sample. We used a four-hour pulse of 4sU immediately before RNA extraction. Based on the findings from Erhard et al. (41), this duration is expected to provide sufficient labelling time to accurately estimate the half-life of genes ranging from approximately one to 10 hours. To increase the detection of a sufficient number of conversions per transcript, we used 500 *µ*M 4sU and sequenced to the libraries 150 bp from both ends (Figure 4a). Our results demonstrated progressive degradation of unlabelled, pre-existing RNA and concurrent transcription of 4sU-labelled, newly synthesised RNA over time (Figure 4g).

PCA was performed on log-transformed normalised read counts. We assessed potential outliers using z-scores calculated from the first two principal components (PC1 and PC2), applying a threshold of |*z*| *>* 3. One sample from the young female group was identified as an outlier due to its significant deviation in PCA space and was subsequently excluded from further analysis (Supplemental Figure 2).

### Ageing has subtle global impact on RNA stability but alters turnover of specific transcripts in articular chondrocytes

grandR was used to estimate both RNA half-lives and synthesis rates for each gene by fitting a standard kinetic expression model to changes in pre-existing and newly synthesised RNA over the progressive labelling time (Supplementary Table 7) (42). Using the raw new-to-total (NTR) RNA values, PCA separated young and old samples into distinct groups along PC1 (24.4% variance) and PC2 (17.7% variance) (Figure 4b). Despite this separation, the mean RNA half-life exhibited minimal changes between age groups (Figure 4c), and the distribution of transcript half-lives within the 1–10h range remained consistent with age (Figure 4d). Empirical cumulative distribution function (ECDF) analysis further confirmed this modest change, with the curve for the old samples indicating a slight overall shift toward shorter RNA half-lives (Figure 4e). Similar to half-life, RNA synthesis rate, regulated by both RNA processing and transcription, was comparable between age groups (Supplementary Figure 3c-e).

Whilst the data indicated that the effect of ageing on transcript half-life in articular chondrocytes is subtle, we observed examples of altered mRNA stability in cells from old horses compared to young. We found a total of 952 genes with half-lives between one and ten hours that exhibited at least a 25% change in stability with age (Supplementary Table 8). GO analysis of this gene set identified enrichment in genes associated with processes such as ribosomal RNA transcription, vitamin metabolism, protein/nucleic acid modification and organelle assembly (Figure 4f). Assessment of PLIN3, which showed loss of a distal polyadenylation site in aged cells, revealed reduced overall transcript levels. However, its half-life was not decreased to the same extent, suggesting that changes in transcript turnover may be associated with reduced usage of the distal polyadenylation site (Figure 4g).

### Integration of RNA expression and stability analysis identified age-related transcriptomic changes

To explore age-related changes in gene expression, we used grandR, which enabled robust quality control and kinetic modelling (42). We identified differentially expressed genes between young and old samples (Supplementary Table 9). PCA using the raw RNA count demonstrated a clearer separation between young and old samples along PC1 (51% variance) and PC2 (19% variance) (Figure 5a), compared to PCA performed with NTR values (Figure 4b). We identified 177 genes that were upregulated, and 185 genes were downregulated in old samples compared to young (Figure 5b and Supplementary Table 10). These differentially expressed genes were not restricted to a particular expression level but were instead found across the full range of expression values (Figure 5c). A heatmap of these differentially expressed genes revealed a distinct expression pattern between young and old samples (Figure 5d). KEGG pathway analysis of this gene set identified enrichment in processes such as translation initiation and elongation, ribosome homeostasis, and WT1/Wnt signalling (Figure 5e). Consistently, GO analysis revealed significant age-related regulation of genes involved in ribosome function and glycoprotein processing (Figure 5f). Taken together with the APA changes we observe in this study, these results suggest that age-associated shifts in polyadenylation may modulate the stability and translational efficiency of key mRNAs, thereby contributing to the broad changes in ribosome function and signalling pathways seen in old cartilage samples.

**Fig. 5.**
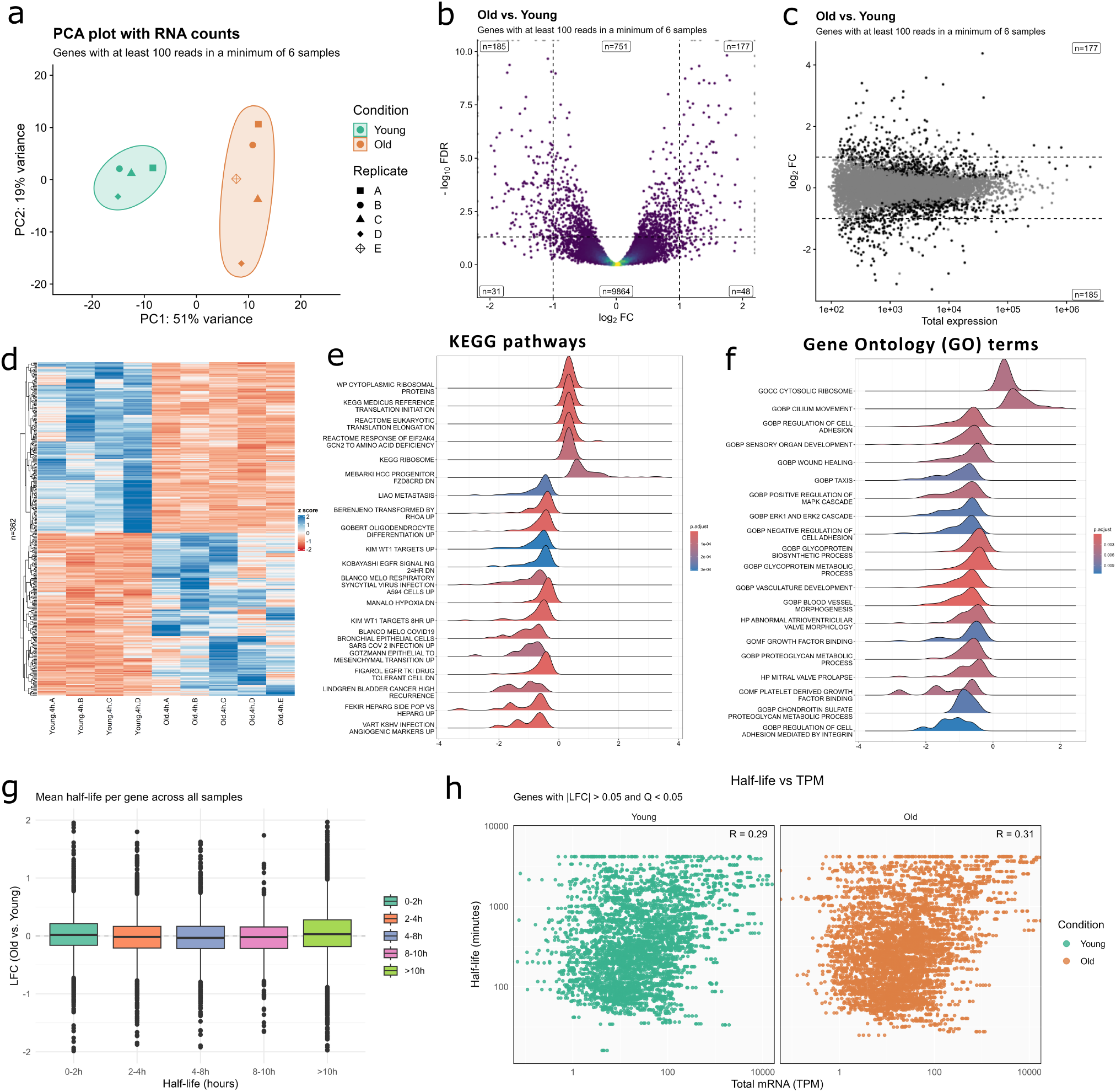
Integration of RNA expression and stability analysis identified age-related transcriptomic changes. (a) PCA analysis with RNA counts. Genes included had *≥* 100 reads in at least 6 samples. (b) Volcano plot illustrates log_2_ fold changes (x-axis) versus *−* log_10_(FDR) for genes with at least 100 reads in a minimum of 6 samples. (c) MA plot of genes with *≥* 100 reads in *≥* 6 samples. Log_2_ fold change is plotted against log_10_ total expression. Black points indicate significantly differentially expressed genes; grey points are non-significant. (d) Heatmap showing the expression levels of significantly differentially expressed genes (|*LFC*| *>* 1) across young and old samples. The colour code scale indicates the normalised counts (ranging from −2 (blue) up to 2 (red)). (e) Ridge plots of Gene Set Enrichment Analysis (GSEA) results for the top 20 KEGG pathways in Equus caballus, comparing old vs. young samples. Colour indicates adjusted p-values, with red representing the most significant enrichment. (f) Ridge plots of GSEA results for the top Gene Ontology (GO) terms in Equus caballus, comparing old vs. young samples. Colour indicates adjusted p-values, with red representing the most significant enrichment. (g) Box plots demonstrating log_2_ fold change (old vs. young) in gene expression across half-life categories. Genes are grouped by half-life ranges (0-2h, 2-4h, 4-8h, 8-10h, *>*10h). Each box represents the median and interquartile range; outliers are shown as individual points. (h) Scatter plots of total RNA levels (TPM) versus RNA half-lives (minutes) in young and old samples. Each dot represents a gene with |*LFC*| *>* 0.5 and Q *<* 0.05. Pearson correlation coefficients are R = 0.29 (young) and R = 0.31 (old).

We compared LFC (old vs. young) across genes stratified by RNA half-life; our results exhibited no systematic shift in LFC median with increasing transcript stability (Figure 5g). Instead, median values across all half-life bins were close to zero, indicating that ageing does not systematically change gene expression as a function of transcript stability. However, the variability of LFC increased with RNA half-life, with long-lived RNAs (*>*10 h) exhibiting greater transcriptional divergence in response to ageing (Figure 5g). We next compared RNA halflives of genes with |*LFC*| *>* 0.5 and Q *<* 0.05 against transcript per million (TPM) values to investigate the contribution of RNA stability to transcript abundance. Across both young and old samples, we observed a modest positive correlation between half-life and TPM (R = 0.29 for young, R = 0.31 for old), suggesting that significantly differentially expressed transcripts with longer half-lives tend to accumulate higher levels (Figure 5h). Notably, the contribution of RNA stability to transcript abundance remained largely unchanged with age. However, these findings suggest that RNA half-life alone does not fully explain steady-state transcript levels in differentially expressed genes with ageing. Consistently, we found a much stronger correlation between synthesis rate and TPM in both young and old samples (R=0.63 for both groups, Supplementary Figure 3e). This indicates that the synthesis rate is the dominant determinant of steady-state transcript abundance in our ageing dataset (Supplementary Figure 3a and 3e).

A recent genome-wide association study (GWAS) meta-analysis of human OA, involving 1,962,069 individuals, identified over 300 gene-associated polymorphisms that confer susceptibility to disease (46). We compared our transcriptomic data with genes identified through GWAS and found that 22 of these genes were differentially expressed in aged samples. Additionally, 36 genes exhibited changes in RNA half-life of at least 25%. Notably, eight genes, vestigial-like family member 4 (VGLL4), stem-loop binding protein (SLBP), AXL receptor tyrosine kinase (AXL), piezo-type mechanosensitive ion channel component 1 (PIEZO1), WSC domain containing 2 (WSCD2), carbohydrate sulfotransferase 3 (CHST3), calcyclin binding protein (CACYBP), and fibroblast growth factor 18 (FGF18), were both differentially expressed and exhibited RNA half-life changes exceeding 25% with age (Supplementary Table 11).

### Age-driven changes in polyadenylation site usage and mRNA half-life in genes with significant differential expression in cartilage

It is well-established that APA events have a great influence on RNA stability (47). To explore this relationship, we filtered differentially expressed genes that exhibited a *≥* 20% change in RNA half-life in our SLAM-seq data and compared them with genes that demonstrated a *≥* 20% shift in APA site in our QuantSeq data. This identified five genes: polypyrimidine tract binding protein 2 (PTBP2), glucosaminyl N-acetyl transferase 4 (GCNT4), xylosyltransferase 1 (XYLT1), Tyrosine-protein phosphatase non-receptor type 2 (PTPN2), and PLIN3 (Figure 6). PTBP2 (Figure 6b) and PTPN2 (Figure 6a). Among the significantly differentially used polyadenylation sites in PTBP2, the APA site at chr5: 65,635,343-65,635,544, located in the 3′UTR, exhibited the most pronounced change in old samples, with over 25% increase in usage (Figure 6b). Notably, PTBP2 had two APA sites in the 3′UTR used differently between old and young samples (Figure 6e). GCNT4, a gene that plays a role in altered glycosylation patterns in cartilage, had a 28.9% shorter RNA half-life in old samples. These genes had two significantly different APA sites located in the 3′UTR (Figure 6c). XYLT1, which plays a critical role in the biosynthesis of glycosaminoglycans (GAGs), demonstrated a significant down-regulation in gene expression and half-life in old cartilage. The APA site at chr13:29,769,639–29,769,846, located in the 3′UTR of XYLT1, exhibited the most pronounced change in old samples (Figure 6d). These data suggest that APA site usage is altered during ageing, and such APA events correlate with changes in RNA half-life, which might affect the stability of gene expression in key age-associated genes.

**Fig. 6.**
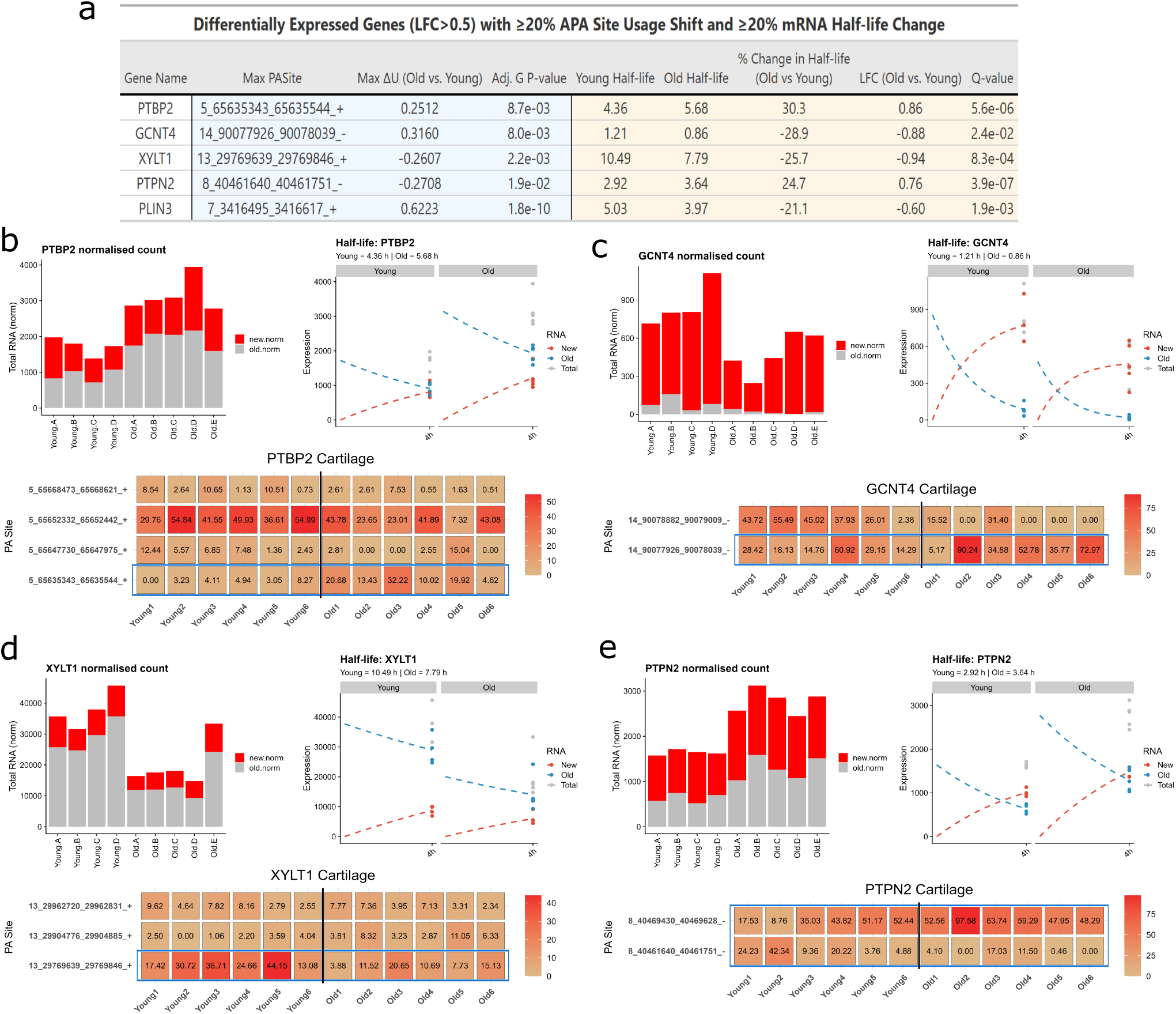
Age-driven changes in polyadenylation site usage and mRNA half-life in genes with significant differential expression in cartilage. (a) The table shows differentially expressed genes with *≥* 20%alternative polyadenylation (APA) site usage shift and *≥* 20% mRNA half-life change. Genes shown were identified by cross-comparing results from QuantSeq (for APA analysis, old vs. young), highlighted in blue, and 4sU-labelled RNA analysed with the grandR and GRAND-SLAM pipeline (for mRNA half-life estimation), highlighted in yellow. This filtering yielded five genes: polypyrimidine tract binding protein 2 (PTBP2), glucosaminyl N-acetyl transferase 4 (GCNT4), xylosyltransferase 1 (XYLT1), Tyrosine-protein phosphatase non-receptor type 2 (PTPN2), and PLIN3. (b-e) Normalised total RNA counts for (b) PTBP2, (c) GCNT4, (d) XYLT1, and (e) PTPN2 in young and old samples. Bars represent newly synthesised (red) and pre-existing (grey) RNA components. (b-e) Progressive labelling plots showing RNA expression dynamics of each gene in young and old cartilage. Plots show newly synthesised (red dashed) and pre-existing (blue dashed), and total (grey dots) RNA, with estimated mRNA half-lives. (b-e) Normalised APA usage matrices for each gene, highlighting the most differentially used polyadenylation site in old vs. young samples (annotated in blue). n = 6.

### Sex exerts a greater influence on RNA stability than age

We also investigated the effects of sex on RNA kinetics in horse chondrocytes and observed substantial differences in RNA half-life between cells from young female and male samples (Supplementary Table 12). PCA of NTR values separated young female and male samples into distinct clusters along PC1 (35.9% variance) and PC2 (16.2% variance) (Figure 7a). The mean RNA half-life differed markedly between groups (Figure 7b); the regression line had a slope of 1.25 (y =1.25x), indicating that RNA transcripts in young males were, on average, 25% more stable than those in young females. Consistent with this finding, the distribution of transcript half-lives within the 1–10h range exhibited a distinct sex-specific pattern (Figure 7c). ECDF analysis further confirmed this difference, with the curve for the young female samples demonstrating a pronounced shift toward shorter RNA half-lives beginning from transcripts with a half-life of two hours or longer (Figure 7d). In parallel with RNA half-life, RNA synthesis rate also showed a distinct sex-specific pattern (Supplementary Figure 4a and 4b), with young males exhibiting a slower synthesis rate. Even though the relationship between RNA half-life and synthesis rate is not strictly inverse, our data from young female and male samples align with the general model: more stable RNAs require less frequent synthesis to maintain transcript levels. Together, these results show clear sex-specific differences in RNA kinetics in horse chondrocytes: young males demonstrate globally longer RNA half-lives and slower synthesis rates compared to young females, consistent with a coordinated regulation of transcript stability and production.

**Fig. 7.**
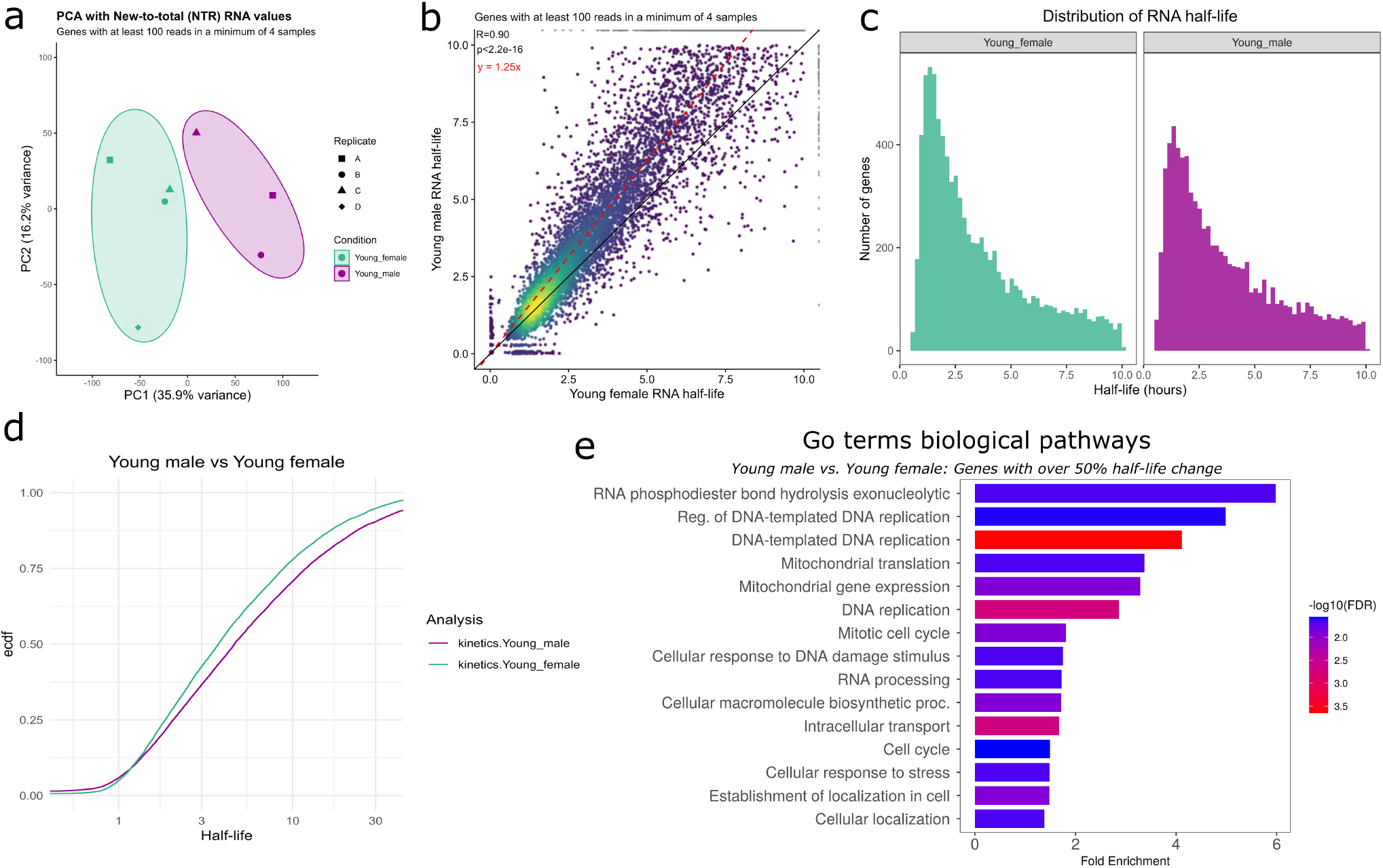
GRAND-SLAM and grandR pipeline reveals longer RNA half-lives in young male cartilage compared to young females. (a) PCA analysis with new-to-total (NTR) RNA values. Genes included had *≥* 100 reads in at least 4 samples. (b) Scatter plot showing the relationship between RNA half-lives in young male and female cartilage for genes with *≥* 100 reads in *≥* 4 samples. A linear regression line (black) is fitted to the data, alongside the reference line y=1.25x (red), indicating substantial differences. Point density is visualised using a colour gradient from blue (low density) to yellow (high density). (c) Distribution of RNA half-lives measured in chondrocytes from young female and male horses. (d) Empirical cumulative distribution function (ECDF) of RNA half-lives in chondrocytes from young female and male horses. The ECDF shows the proportion of genes with half-lives below a given threshold, highlighting longer RNA half-lives in young males compared to females. (e) Gene Ontology (GO) biological process enrichment analysis of genes exhibiting *>* 50% change in RNA half-life in young male cartilage samples compared to young female. Bar length indicates fold enrichment, and bar colour represents *−* log_10_(FDR)values associated with pathway enrichment, ranging from blue (lower significance) to red (higher significance).

To determine the potential functional impact of the sex-linked difference in mRNA half-life, we filtered for genes with half-lives between one and ten hours that exhibited at least a 50% change in RNA stability in young male samples compared to females, with 783 genes meeting this threshold (Supplementary Table 13). GO enrichment analysis identified significant overrepresentation of key cellular processes related to regulation of DNA replication, mitochondrial translation, mitochondrial gene expression, mitotic cell cycle, and cellular response to DNA damage (Figure 7e).

### Sex-dependent RNA stability influences transcript abundance in chondrocytes

We next identified differentially expressed genes in young males compared to young females (Supplementary Table 14). PCA of RNA counts exhibited a clear separation between the groups along PC1 (53% variance) and PC2 (20% variance) (Figure 8a). We identified a total of 334 genes that were differentially regulated in male samples compared to females (Figure 8b and supplementary Table 15). These differentially expressed genes were found across the full range of expression values (Figure 8c). A heatmap of these differentially expressed genes revealed a distinct expression pattern between female and male samples (Figure 8d). Using these differentially expressed genes, we performed gene set enrichment analysis with hallmark gene sets. This identified epithelial–mesenchymal transition, angiogenesis, apical junction, coagulation, and myogenesis pathways demonstrated significant enrichment. Notably, stress response and several immune-related pathways, including TNF*α* signalling via NF-*κ*B, L6/JAK/STAT3 signalling, interferon gamma response, and inflammatory response, exhibited skewed expression distributions and significant enrichment.

**Fig. 8.**
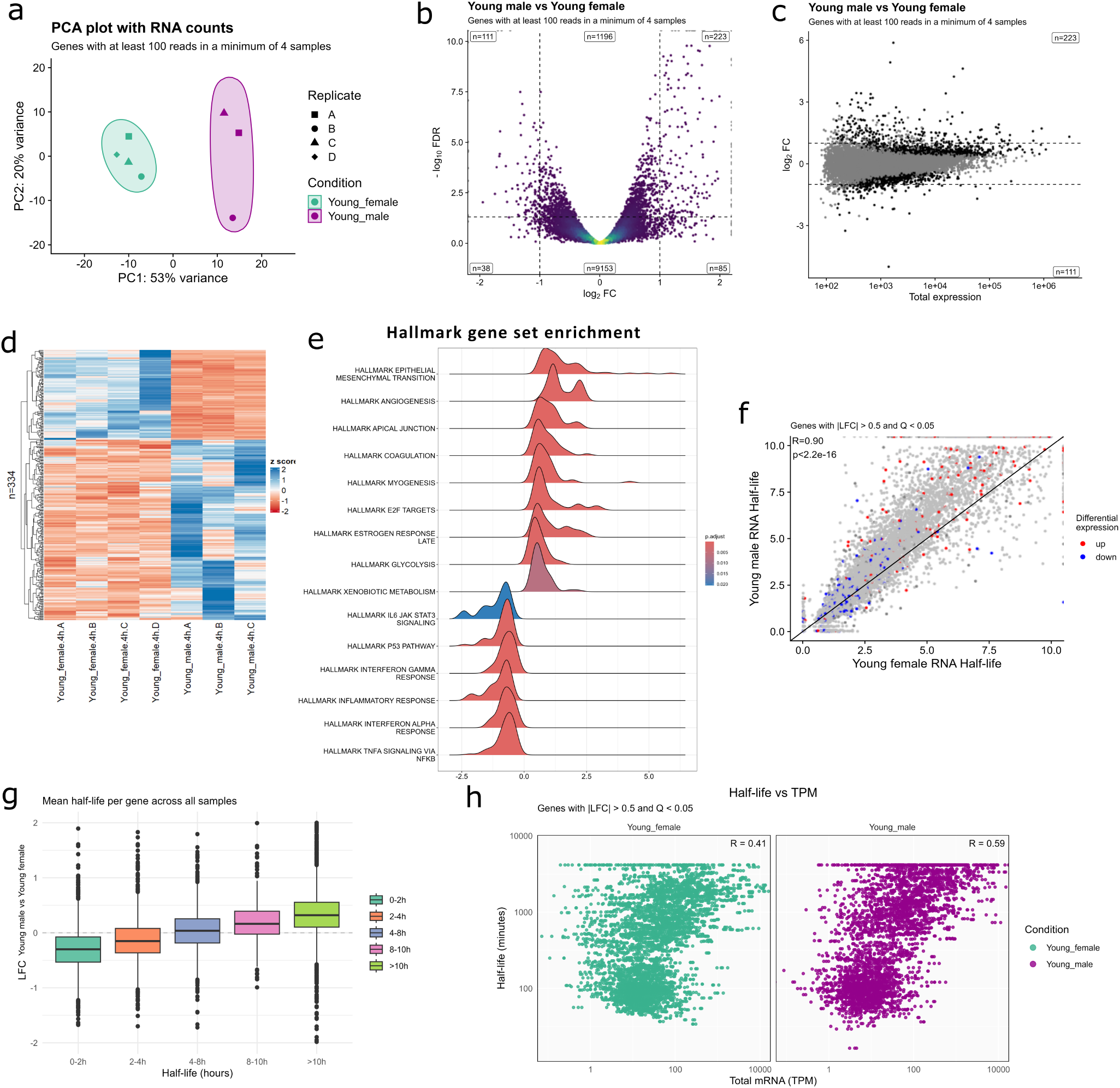
Transcriptome-wide analysis of 4sU-labelled RNA using the grandR pipeline in young male and female cartilage. (a) PCA analysis with RNA counts. Genes included had *≥* 100 reads in at least 4 samples. (b) Volcano plot illustrates log_2_ fold changes (x-axis) versus *−* log_10_(FDR) (y-axis) for genes with at least 100 reads in a minimum of 4 samples. (c) MA plot of genes with *≥* 100 100 reads in *≥* 4 samples. Log_2_ fold change is plotted against log_10_ total expression. Black points indicate significantly differentially expressed genes; grey points are non-significant. (d) Heatmap showing the expression levels of significantly differentially expressed genes (|*LFC*| *>* 1) across young female and male samples. The colour code scale indicates the normalised counts (ranging from −2 (blue) up to 2 (red)). (e) Ridge plots of hallmark gene set enrichment analysis comparing young male and female in Equus caballus cartilage. Colour indicates adjusted p-values, with red representing the most significant enrichment. (f) Scatter plot showing the relationship between RNA half-lives in young male and female cartilage for genes with *≥* 100 reads in *≥* 4 samples. Genes are coloured by differential expression: red for up-regulated, blue for down-regulated (|*LFC*| *>* 0.5, Q *<* 0.05), and grey for non-significant. (g) Box plots showing log_2_ fold change in gene expression (old vs. young) across RNA half-life categories. Genes are grouped by half-life ranges: 0-2h, 2-4h, 4-8h, 8-10h, and *>*10h. Each box represents the median and interquartile range; outliers are shown as individual points. The plot suggests that genes with longer RNA half-lives tend to be upregulated in young male cartilage samples compared to young female. (h) Scatter plots of total RNA levels (TPM) versus RNA half-lives (minutes) in young female and male cartilage samples. Each dot represents a gene with |*LFC*| *>* 0.5 and Q *<* 0.05. Pearson correlation coefficients are R = 0.41 (female) and R = 0.59 (male), indicating a moderate positive relationship between RNA abundance and stability, with a stronger correlation observed in males.

To investigate the relationship between sex-specific gene expression and RNA stability, we compared LFC (young male vs. young female) across genes stratified by RNA half-life. Our results showed a systematic shift in median LFC with increasing transcript stability (Figure 8g), indicating a sex-specific systematic change in gene expression as a function of transcript stability. Notably, the long-lived RNAs (*>*10 h) exhibited a greater sex-specific transcriptional divergence (Figure 8g), consistent with patterns observed in our ageing dataset. We next compared RNA half-lives of genes with |*LFC*| *>* 0.5 and Q *<* 0.05 against TPM values to investigate the contribution of RNA half-life to transcript abundance. As shown in Figure 8h, the correlation between transcript stability and abundance was stronger in young males (R=0.59) than in young females (R=0.41), indicating that RNA stability contributes more prominently to determining transcript abundance of differentially expressed genes in males. In contrast, synthesis rate exhibited a stronger correlation with TPM in young females (R=0.51) than in males (R=0.44; Supplementary Figure 4d). This indicates that the synthesis rate is the dominant determinant of steady-state transcript abundance of differentially expressed genes in young females compared to males.

## Discussion

Our findings suggest that fundamental physiological factors, age and sex, exert distinct regulatory pressures on post-transcriptional gene regulation in joint tissues. The observation that ageing alters both polyadenylation site usage and transcript turnover rates in cartilage, even in the absence of overt pathology, supports the hypothesis that molecular ageing processes precede and potentially predispose to degenerative joint disease. Furthermore, the unexpected influence of sex on mRNA turnover in chondrocytes raises the possibility that sex-specific regulatory mechanisms shape RNA stability, potentially contributing to differential susceptibility to joint degeneration between males and females. Together, these results point to age and sex as key drivers of post-transcriptional regulation with potential roles in shaping joint health trajectories.

Using the equine metacarpophalangeal joint as a model system, previously established for studying age-associated changes in the musculoskeletal system (48,49), we have investigated post-transcriptional regulatory mechanisms by comparing tissue from skeletally mature young horses with that from aged individuals. To ensure that OA was not a confounding factor, we applied a well-established equine macroscopic grading system, excluding any joints that showed signs of pathology. We initially employed 3′ end sequencing using the QuantSeq-Reverse technique to identify the usage of polyadenylation sites in young and old horses. Our results demonstrated that APA events are significantly influenced by age in cells from both articular cartilage and synovium, providing additional context to the established effect of age on the transcriptome of joint cells such as chondrocytes (3). To our knowledge, while previous studies have linked APA site usage to specific pathologies, this is the first to directly investigate the impact of mammalian ageing on polyadenylation patterns.

The analysis of polyadenylation events identifies a range of transcript variants linked to ageing (50). Extension or shortening of the 3′UTR is a common consequence of APA. In this study, we observed age-dependent shifts in 3′UTR length in genes such as PEX2 and PLIN3, where chondrocytes from aged individuals expressed transcripts with longer or shorter 3′UTRs, respectively. Similarly, the use of intronic polyadenylation sites can result in truncated protein products. A notable example is an intronic site in the oncostatin M receptor (OSMR) gene, which we previously reported as being regulated in human osteoarthritic cartilage (29), leading to the production of a secreted, antagonistic form of the receptor (51). Our analysis identified numerous intronic polyadenylation sites that showed differential usage between young and aged tissues. Notable examples include sites within the RBFOX1 and PROM1 genes, which were regulated in the synovium. Given that RBFOX1 encodes a splicing factor, these findings suggest a potential link between alternative polyadenylation and other RNA processing events, such as splicing, highlighting a coordinated post-transcriptional regulatory landscape that may be altered with age.

A site in the NRG1 gene that exhibited consistent age-dependent regulation across both cartilage and synovium. NRG1 is particularly interesting, as it has been implicated in cartilage regeneration in zebrafish (52). The NRG1 polyadenylation site we identified corresponds to two equine EST sequences and is located within a large first intron, approximately 100 kb downstream of exon 1, making transcriptional read-through from the upstream exon unlikely. Supporting this, our qRT-PCR analysis detected transcript expression using primers positioned just upstream of the polyadenylation site, but not when primers 461bp upstream of the site were used. These findings suggest that the NRG1-associated polyadenylation event reflects the differential expression of a distinct transcript originating within the first intron. Interestingly, this transcript shows strong homology at its 3′ end with EEF1A1 mRNA and corresponds to the long non-coding RNA NRG1-IT1, which resides within the first intron of the human NRG1 gene. This raises the possibility of an age-associated regulatory mechanism regulating NRG1-IT1 in joint tissues. NRG1 plays a critical role in maintaining cellular structural function (53). Previous studies have shown that NRG1 promotes cell growth, counteracts inflammatory responses, and inhibits apoptosis (54). Ageing substantially affects NRG1 regulation through complex mechanisms that alter its expression, isoform diversity, and signalling activity, particularly in the context of neurodegenerative diseases (55,56). Given the importance of cellular integrity in joint tissues and the established role of NRG1 in cartilage regeneration, the NRG1-associated polyadenylation site identified here may represent a promising therapeutic target in OA.

The age-associated shifts in polyadenylation and transcript turnover we have observed may represent an early molecular mechanism contributing to the progressive functional decline of joint tissues. APA has the potential to alter transcript stability, localisation, and translation efficiency (52), thereby shaping the proteomic landscape of chondrocytes and synoviocytes, whilst transcript turnover rates contribute strongly to how responsive a gene is to regulatory signals (57). In joint tissues, genes involved in cellular integrity and matrix homeostasis, such as collagen and aggrecan, possess long 3′UTR rich in regulatory elements (33,58). APA-driven lengthening of these transcripts during ageing could dampen synthesis of structural or anabolic components, while APA-driven shortening could lead to overproduction of catabolic or inflammatory mediators. Together, these effects provide a plausible mechanism by which age-associated APA shifts could impair the balance between anabolic and catabolic pathways, compromise chondrocyte and synoviocyte function, and ultimately undermine cartilage integrity. As transcript turnover strongly influences responsiveness to regulatory signals, APA-mediated changes in stability may also blunt the ability of joint cells to adapt to mechanical load or injury. Furthermore, we observe distinct polyadenylation patterns between cartilage and synovium, despite sampling paired tissues from the same joint. This design allows us to control for inter-individual variability and highlights the possibility that tissue-specific regulatory pressures shape differential susceptibility within the joint microenvironment, a dimension not addressed in previous studies and one of the key novelties of our work.

Employing SLAM-seq, we analysed transcript half-lives in articular chondrocytes isolated from young and old horses. The chondrocytes were cultured at high density and used within two days of isolation to minimise the effects of culture-based dedifferentiation. SLAM-seq has previously identified pronounced effects on global transcript turnover in other cell types, driven by the knockout of regulatory molecules (57), cellular differentiation (40,59) and stress signals (60). Our data revealed that age also influenced transcript turnover in articular chondrocytes, and we found that nearly 1000 genes exhibited an altered mRNA half-life of 25% or greater in old cells. This indicates that the effects of altered post-transcriptional regulation associated with age are widespread and provides an interesting insight into the functionality of older chondrocytes, where alterations in transcript kinetics across a wide range of genes could potentially lead to differences in cellular responsivity compared to younger cells. Assessment of steady-state gene expression in these samples identified significant changes in expression levels of genes associated with translation initiation and ribosome function, areas that have been highlighted as processes that can contribute to age-associated joint diseases (48,61). We also observed extensive age-related shifts in APA, a mechanism known to affect mRNA stability and translational efficiency. Together, these changes suggest that ageing alters both the protein-synthesis machinery and post-transcriptional mRNA processing in chondrocytes, which could underlie the broad alterations in ribosome function and signalling pathways we detect in old cartilage samples. Interestingly, our comparison of transcript turnover rates with steady-state mRNA expression levels revealed a weak correlation. This suggests that age-associated changes in mRNA decay may be buffered by compensatory transcriptional activity, potentially leading to altered gene responsiveness without substantial changes in overall expression levels.

Our analyses identified that ageing reshapes the cartilage transcriptome through coordinated changes in both transcriptional and post-transcriptional regulation. GO analysis of differentially expressed genes revealed that glycoprotein processing is one of the major age-sensitive pathways. By integrating RNA half-life data with APA profiles, we found that ageing affects not only the expression levels but also the stability and structural configuration of key genes such as XYLT1 and GCNT4. These enzymes represent critical regulatory points in protein glycosylation: XYLT1 establishes the very start of GAG chain assembly, whilst GCNT4 contributes to the structural diversity and complexity of glycoproteins. This is particularly significant given that proteoglycan synthesis is essential for maintaining cartilage integrity and function. Dysregulation of XYLT1 has been implicated in impaired accelerated chondrocyte maturation, extracellular matrix assembly, and early cartilage degeneration, underscoring its potential as both a mechanistic biomarker and therapeutic target in OA pathogenesis (62–64). To our knowledge, while direct evidence linking GCNT4 to OA is limited, it has been identified as one of the most dysregulated genes in the articular cartilage of OA patients (65). Taken together, these results indicate that targeted investigation of APA and mRNA half-life in genes central to glycoprotein metabolism may provide new insights into the molecular underpinnings of OA. Moreover, therapeutic strategies aimed at modulating these APA events could help maintain balanced GAG metabolism and protect against cartilage degeneration in OA.

We were intrigued to discover a pronounced effect of sex on mRNA turnover in articular chondrocytes from young horses, with transcripts in males exhibiting greater stability across the transcriptome. Furthermore, by integrating turnover data with steady-state expression analysis, we found a clear relationship between sex-biased gene expression and transcript stability. Specifically, transcripts expressed at lower levels in males were more likely to be short-lived, suggesting that differential mRNA decay may contribute to sex-specific gene regulation in joint tissues. These results are particularly relevant in the context of OA, where sex is one of the key factors in disease development. Epidemiological data studies consistently show that women have a higher prevalence and severity of OA than men (11,66-68). Our finding that transcript stability is influenced by sex, therefore underlines the importance of investigating sexual dimorphism in cartilage biology. Such differences in post-transcriptional regulation may affect how male and female cells respond to stress or maintain cartilage homeostasis, providing a potential molecular basis for sex-specific variation in joint disease. However, several important considerations must be taken into account when interpreting these data. All male samples were obtained from geldings, meaning systemic testosterone levels would have been negligible from approximately 6–12 months of age (69). It is therefore plausible that some of the observed sex differences may be attributable to the effects of reduced testosterone. Additionally, while female sex is a known risk factor for OA in humans, the influence of sex on OA progression varies across species. For example, in murine models of induced OA, males often exhibit more severe disease than females (70). In horses, to our knowledge, no studies have yet reported sex-specific effects in OA. Overall, our data represent an extensive transcriptome-wide identification of sex-dependent post-transcriptional changes in equine cartilage. These findings offer a novel perspective on the molecular mechanisms underlying sex differences in joint disease and underscore the need for further investigation in human systems. While species differences must be carefully considered, our results provide a compelling rationale for exploring sex-specific regulation as a contributing factor to OA pathogenesis.

This study represents the first comprehensive assessment of the effect of ageing and sex on post-transcriptional gene regulation in a mammalian system. We demonstrate that there are significant effects of age on these processes, but they remain more modestly regulated than in some other model systems driven by differentiation, disease or mutagenesis. In contrast, sex appears to have a greater influence on transcript processing and stability. These insights into mRNA kinetics and polyadenylation pattern provide a critical framework for understanding how ageing and sex shape post-transcriptional regulation in joint tissues.

## Data availability

High-throughput sequencing data (QuantSeq and SLAM-seq) generated for this study have been deposited in the GEO database under accession numbers GSE306645 and GSE306550, respectively. Data from SLAM-seq analysis are available as an interactive resource and for download under: https://velorna.liverpool.ac.uk/AgeingApp/ and https://velorna.liverpool.ac.uk/SexApp/.

## Code availability

PolyA-miner (35), GRAND-SLAM (41), and grandR (42) were used to analyse the datasets. The code underlying these analyses is available online and can be shared upon reasonable request to the corresponding author.

## Supplementary data

Supplementary Data are available online.

## Acknowledgments

We thank all members of the Tew and Peffers laboratories for their help with sample collection and resource management. We would also like to thank the Centre for Genomic Research at the University of Liverpool for providing sequencing services. Finally, we extend our thanks to F. Drury and Sons, Swindon, UK, abattoir for supplying tissue samples.

## Author contributions statement

U.E. performed the experimental work, conducted data curation, formal analysis, and investigation; contributed to visualisation; and was responsible for writing the original draft as well as reviewing and editing the manuscript. S. R. T. was responsible for conceptualisation, methodology, formal analysis, writing the original draft, reviewing and editing the manuscript. S.H., I.K., J. A., and M. J. P. contributed to data curation, formal analysis, software implementation, and resource management. All authors reviewed and approved the final manuscript.

