## Supplementary figure 1 for "Characterising the effect of age and sex on post-transcriptional regulation in synovial joint tissues"

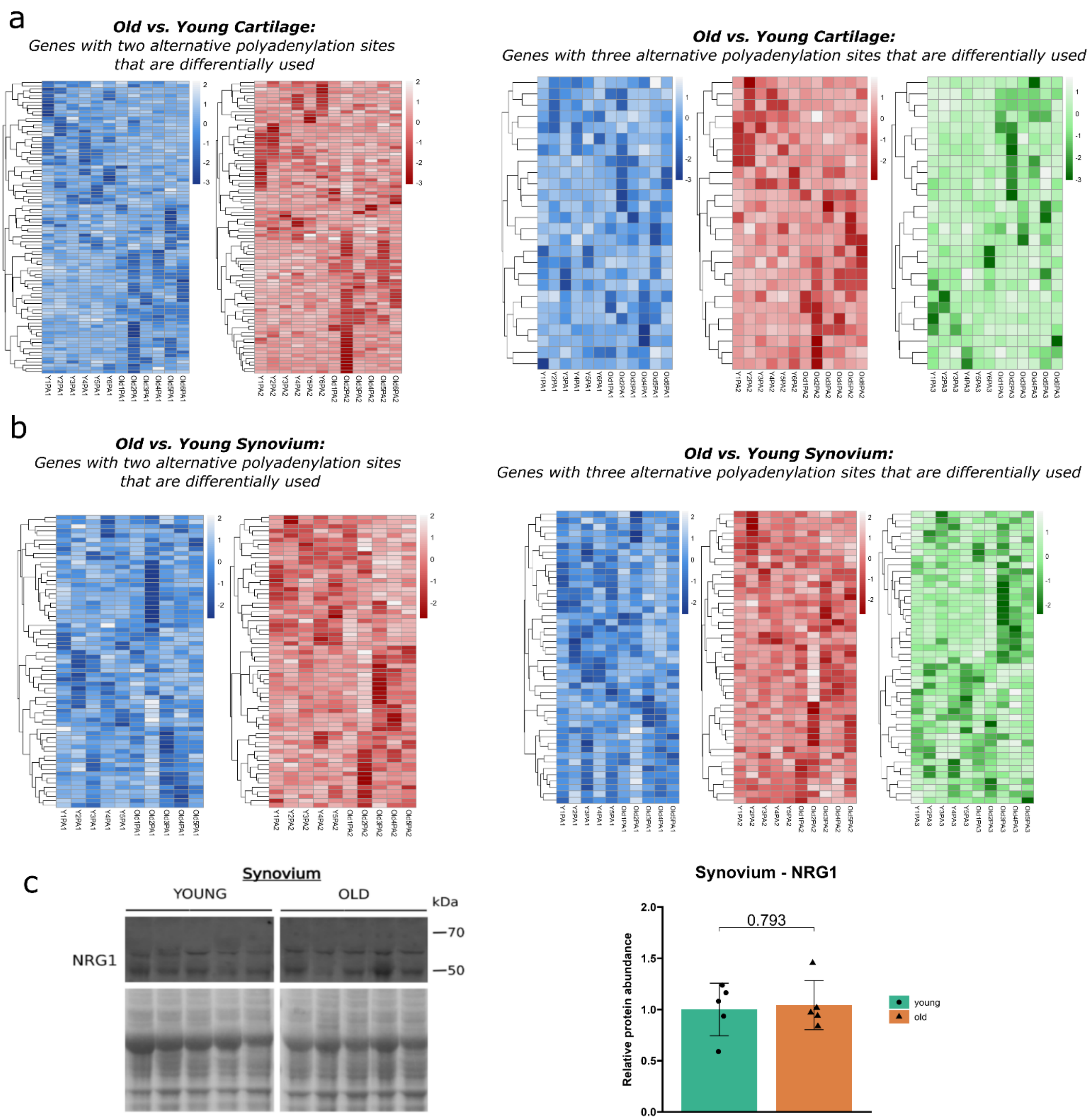


**Supplementary Figure 1. Significant shifts in alternative polyadenylation (APA) site usage in cartilage and synovium from young and old horses and NRG1 protein abundance. (a)** APA heatmap of genes with two polyadenylations and three polyadenylation sites in cartilage. **(b)** APA heatmap of genes with two polyadenylations and three polyadenylation sites in synovium. **(c)** Western blot analysis and quantification of neurogulin1 (NRG1) in synovium. Samples were run on the same gel, and images were cropped solely for presentation purposes. Ponceau staining was used as a loading control.

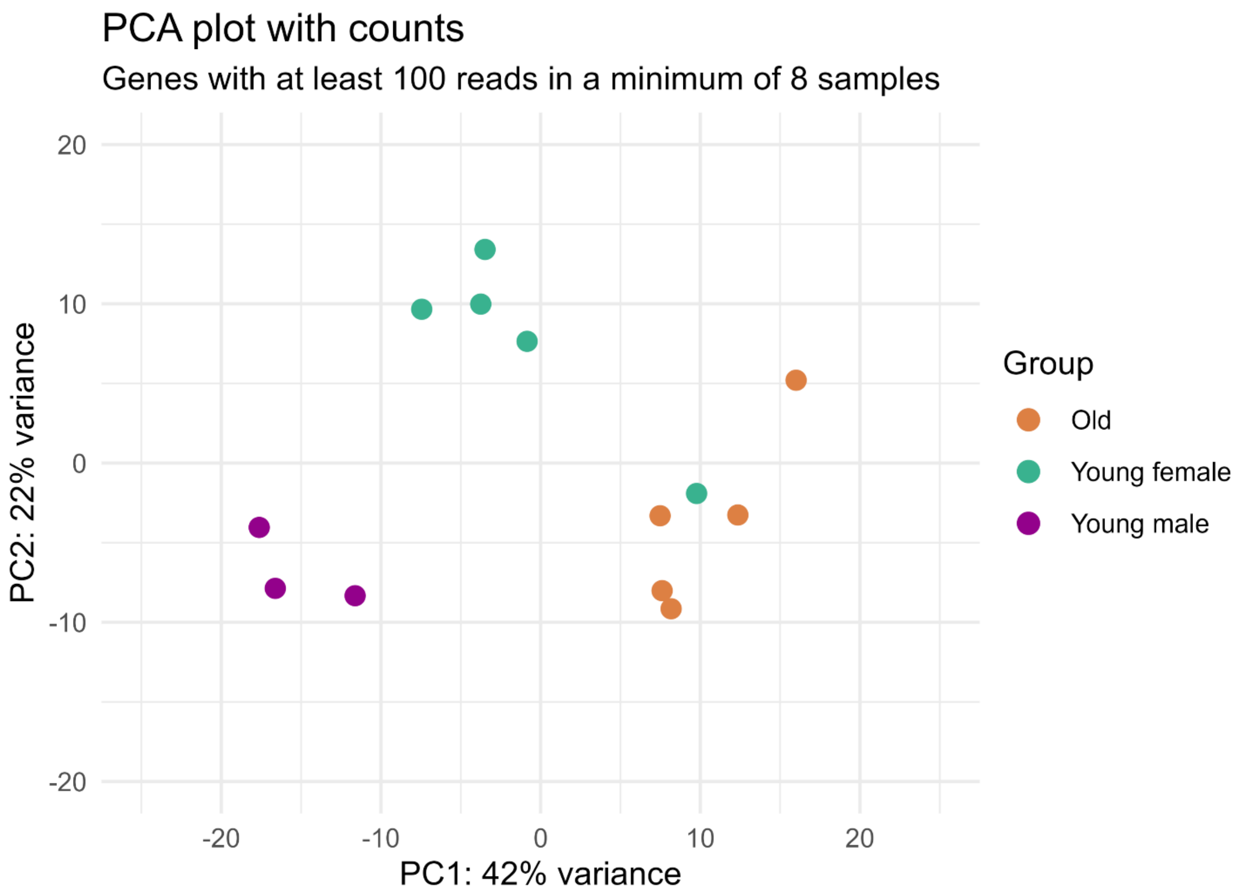


**Supplementary Figure 2. Principal Component Analysis (PCA) of log-transformed normalised read counts across age and sex groups.** PCA was performed on biological replicates from young female, young male, and old groups. Each point represents an individual sample, colour-coded by group: green (young female), purple (young male), and orange (old). The first two principal components (PC1 and PC2) explain 42% and 22% of the total variance, respectively. Outlier detection was conducted using z-scores derived from PC1 and PC2 coordinates, with a threshold of $|z| > 3$. One from the young female group was identified as an outlier due to its significant deviation in PCA space.

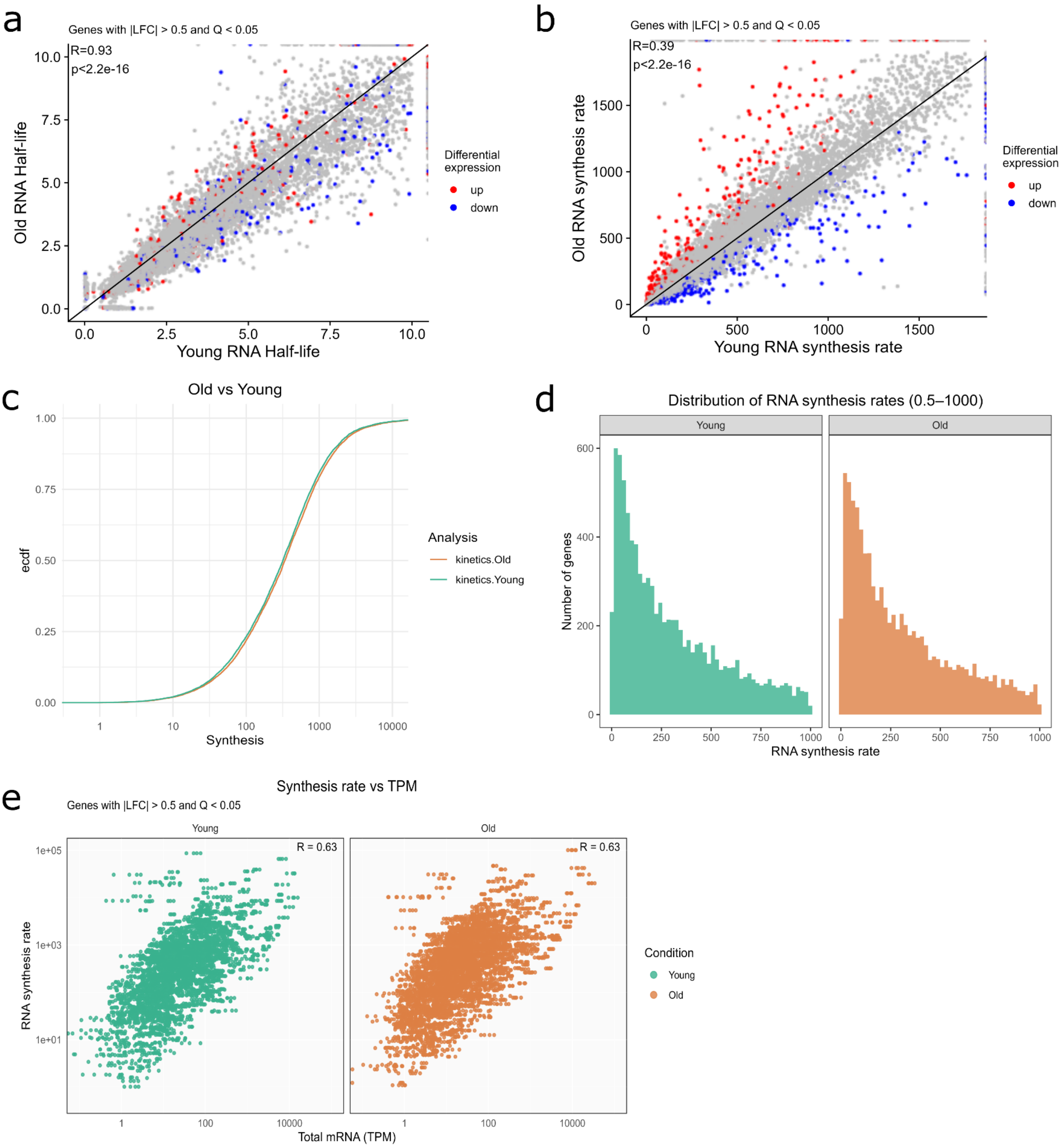


**Supplementary Figure 3. RNA half-life and synthesis rates in young and old samples.** Scatter plot showing the relationship between RNA half-lives **(a)** and synthesis rates **(b)** in young and old samples for genes with ≥100 reads in ≥6 samples. Genes are coloured by differential expression: red for up-regulated, blue for down-regulated (|LFC| > 0.5, Q < 0.05), and grey for non-significant. **(c)** Empirical cumulative distribution function (ECDF) of RNA synthesis rate for chondrocytes from young and old horses. The ECDF illustrates the proportion of genes with half-lives below a given threshold across both age groups. **(d)** Distribution of RNA synthesis rate measured in chondrocytes from young and old samples by SLAM-seq. **(e)** Scatter plots of total RNA levels (TPM) versus RNA synthesis rate (minutes) in young and old samples. Each dot represents a gene with |LFC| > 0.5 and Q < 0.05. Pearson correlation coefficients are R = 0.63 for both groups.

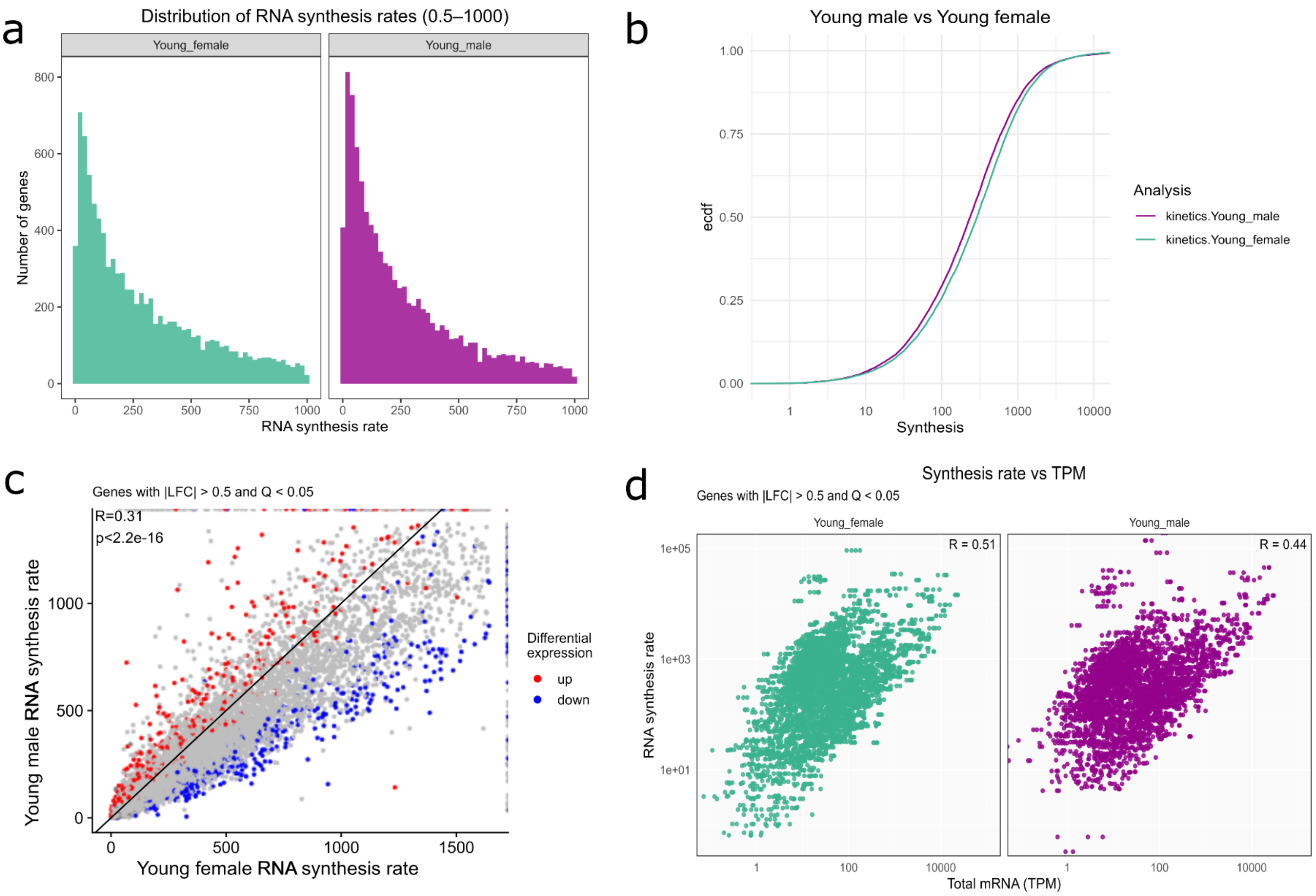


**Supplementary Figure 4 RNA half-life and synthesis rates in young female and young male samples. (a)** Distribution of RNA synthesis rate measured in chondrocytes by SLAM-seq. **(c)** Empirical cumulative distribution function (ECDF) of RNA synthesis rate for chondrocytes from female and male samples. **(b)** Empirical cumulative distribution function (ECDF) of RNA synthesis rate for chondrocytes, indicating a slower RNA synthesis rate in young males compared to females. **(c)** Scatter plot showing the relationship between RNA synthesis rates. Genes are coloured by differential expression: red for up-regulated, blue for down-regulated (|LFC| > 0.5, Q < 0.05), and grey for non-significant. **(d)** Scatter plots of total RNA levels (TPM) versus RNA synthesis rate (minutes) in young female and male samples. Each dot represents a gene with |LFC| > 0.5 and Q < 0.05. Pearson correlation coefficients are R = 0.51 for females and R=0.44 for males.
